# Nonquantal Transmission at the Vestibular Hair Cell-Calyx Synapse: K_LV_ Currents Modulate Fast Electrical and Slow K^+^ Potentials in the Synaptic Cleft

**DOI:** 10.1101/2021.11.18.469197

**Authors:** Aravind Chenrayan Govindaraju, Imran H. Quraishi, Anna Lysakowski, Ruth Anne Eatock, Robert M. Raphael

**Author notes:** Author for correspondence: Robert M. Raphael, Dept. of Bioengineering, MS 142, Rice University, Houston, TX.

## Abstract

Vestibular hair cells transmit information about head position and motion across synapses to primary afferent neurons. At some of these synapses, the afferent neuron envelopes the hair cell, forming an enlarged synaptic terminal called a calyx. The vestibular hair cell-calyx synapse supports a mysterious form of electrical transmission that does not involve gap junctions termed nonquantal transmission (NQT). The NQT mechanism is thought to involve the flow of ions from the pre-synaptic hair cell to the post-synaptic calyx through low-voltage-activated channels driven by changes in cleft [K^+^] as K^+^ exits the hair cell. However, this hypothesis has not been tested with a quantitative model and the possible role of an electrical potential in the cleft has remained speculative. Here we present a computational model that captures salient experimental observations of NQT and identifies overlooked features that corroborate the existence of an electrical potential (*ϕ*) in the synaptic cleft. We show that changes in cleft *ϕ* reduce transmission latency and illustrate the relative contributions of both cleft [K^+^] and *ϕ* to the gain and phase of NQT. We further demonstrate that the magnitude and speed of NQT depend on calyx morphology and that increasing calyx height reduces action potential latency in the calyx afferent. These predictions are consistent with the idea that the calyx evolved to enhance NQT and speed up vestibular signals that drive neural circuits controlling gaze, balance, and orientation.

**Significance Statement:** The ability of the vestibular system to drive the fastest reflexes in the nervous system depends on rapid transmission of mechanosensory signals at vestibular hair cell synapses. In mammals and other amniotes, afferent neurons form unusually large calyx terminals on certain hair cells, and communication at these synapses includes nonquantal transmission (NQT), which avoids the synaptic delay of quantal transmission. We present a quantitative model that shows how NQT depends on the extent of the calyx covering the hair cell and attributes the short latency of NQT to changes in synaptic cleft electrical potential caused by current flowing through open potassium channels in the hair cell. This previously undescribed mechanism may act at other synapses.

## Introduction

The vestibular inner ear transduces head motion and drives the fastest known reflexes in the nervous system, including the vestibulo-ocular reflex (Huterer and Cullen 2002) and the vestibulo-collic reflex (Colebatch and Rosengren 2019). These reflexes support gaze, balance, and orientation (Cullen 2019) and are essential for locomotion. The first synapses in these pathways are from mechanosensory hair cells onto primary afferent nerve terminals. In mammals and other amniotes, many vestibular afferent terminals form prominent cup-like structures, termed calyces around specialized type I hair cells (Wersäll 1956; Gulley and Bagger-Sjöbäck 1979; Lysakowski and Goldberg 2004). Transmission at these synapses has been inferred to be exceptionally fast (McCue and Guinan 1994), prompting speculation that it includes a direct, electrical component in addition to glutamate exocytosis from synaptic vesicles (quantal transmission). Electrical transmission typically involves gap junctions (Nielsen et al. 2012) but gap junctions are not present at vestibular hair cell-calyx (VHCC) synapses (Gulley and Bagger-Sjöbäck 1979; Ginzberg 1984; Yamashita and Ohmori 1991). Nonetheless, electrophysiological recordings from calyx-bearing afferents established that an additional mode of transmission can occur with or without quantal transmission (Yamashita and Ohmori 1990; Holt et al. 2007; Songer and Eatock 2013; Highstein et al. 2014; Contini et al. 2017). This mysterious form of transmission came to be called Nonquantal Transmission (NQT).

NQT has been suggested to be mediated by the flow of ions from the pre-synaptic hair cell to the post-synaptic calyx through low-voltage-activated K^+^ (K_LV_) channels (Eatock 2018), but how pre-synaptic currents alter the driving forces of post-synaptic currents has not been well understood. A key challenge for understanding NQT at the VHCC synapse is the lack of experimental access to the synaptic cleft and pre-and post-synaptic membranes. Early models (Chen 1995; Goldberg 1996) recognized that the elongated cleft space between the type I hair cell and enveloping calyx structure could provide increased electrical resistance and limit the diffusion of ions. In these formulations, K^+^ that enters hair cells during mechanotransduction is extruded into the synaptic cleft, leading to K^+^ accumulation and a change in the K^+^ chemical potential that drives membrane currents. Changes in cleft electrical potential were not thought to be significant. In support of these models, subsequent experimental results emphasized the role of K^+^ accumulation (Lim et al. 2011; Contini et al. 2012, 2017, 2020; Spaiardi et al. 2020). Other processes were also suggested, including accumulation of glutamate (Dalet et al. 2012; Sadeghi et al. 2014) and protons (Highstein et al. 2014). However, the existence and role of an electrical potential in the synaptic cleft has remained a matter of speculation. A clear biophysical explanation for how NQT occurs and how currents and driving forces interact quantitatively at the VHCC synapse has not yet been presented.

To discern the biophysical basis of NQT, we have developed a detailed computational model of the VHCC synapse based on the anatomy of the afferent calyx terminal and the properties of ion channels and transporters expressed in the hair cell, calyx terminal and afferent fiber. Simulated responses of the VHCC model to hair bundle deflection and voltage-clamp steps accurately predict experimental data. Such validation gives us confidence in the model’s predictions for sites with no experimental data, notably the inaccessible synaptic cleft. Results show that NQT at the VHCC synapse depends on the geometry of the calyx, involves modulation of both the electrical potential and K^+^ concentration in the synaptic cleft, and is mediated primarily by currents through low-voltage-activated potassium (K_LV_) conductances that take different forms in the presynaptic and postsynaptic membranes (“g_K,L_” channels and K_V_7 channels, respectively). A major outcome of the model is the recognition of features in post-synaptic responses (Songer and Eatock 2013; Contini et al. 2020) as evidence for changes in electrical potential in the synaptic cleft during NQT.

Changes in extracellular electrical potential have long been known to play a role in ephaptic coupling, a general term for coupling via proximity of neural elements (Jefferys 1995; Faber and Pereda 2018). Ephaptic coupling has been implicated in signal transmission in the retina (Vroman et al. 2013; Warren et al. 2016), cerebellum (Han et al. 2018, 2020), hippocampus (Chiang et al. 2019), and heart (Veeraraghavan et al. 2014; Gourdie 2019). At the VHCC synapse, however, the importance of changes in cleft electrical potential have gone unrecognized despite the extensive synaptic cleft which is formed by the close apposition of the pre-synaptic (hair cell) and post-synaptic (afferent) membranes. The VHCC model we develop is able to decouple electrical and K^+^ potentials and demonstrate that change in cleft electrical potential is necessary to explain: 1) the short latency of NQT; 2) the experimentally observed phase response of NQT (Songer and Eatock 2013); 3) fast retrograde events in the hair cell during post-synaptic action potentials (Contini et al. 2017); and 4) the shape of the fast current response of the calyx to voltage changes in the hair cell (Contini et al. 2020). By showing how the magnitude and speed of NQT depends on the calyx, our model supports the idea that calyx evolved to support faster transmission in response to increased locomotory challenges presented by the tetrapod transition from water to land (Eatock 2018; Curthoys et al. 2019).

### Key Abbreviations

AP: action potential
MET: mechanoelectrical transduction
NQT: Nonquantal Transmission
VHCC: Vestibular Hair Cell-Calyx

### Electrical potentials

(with respect to perilymph):

*ϕ_H_*: electric potential in the hair cell
*ϕ_SC_*: electrical potential in the synaptic cleft
*ϕ_C_*: electric potential in calyx

### Transmembrane Voltages

*V_H_*: voltage across the basolateral hair cell membrane, (*ϕ_H_* – *ϕ_SC_*)
*V_CIF_*: voltage across calyx inner face membrane, (*ϕ_C_* – *ϕ_SC_*)

### Concentrations and ionic potentials

[*K*^+^]_*SC*_: ^K+^ concentration in the synaptic cleft
[*Na*^+^]_*SC*_: Na^+^ concentration in the synaptic cleft
*E_κ_*: Equilibrium potential for K^+^ (K^+^ potential) across hair cell or calyx membranes

## General Description of the Model

The model has been developed for an afferent calyx around a single type I hair cell (“simple calyx-only terminal”) based on the geometry of a vestibular hair cell and calyx from the striolar zone of the rat utricle, see **Fig. 1** and **Supplementary Information (*SI*), Fig. S1**. The morphology of vestibular hair cells and calyces is highly conserved within particular zones of vestibular epithelia (Lysakowski and Goldberg 1997, 2008) and thus the model is representative of simple VHCC synapses in the striolar zone.

**Fig. 1.**
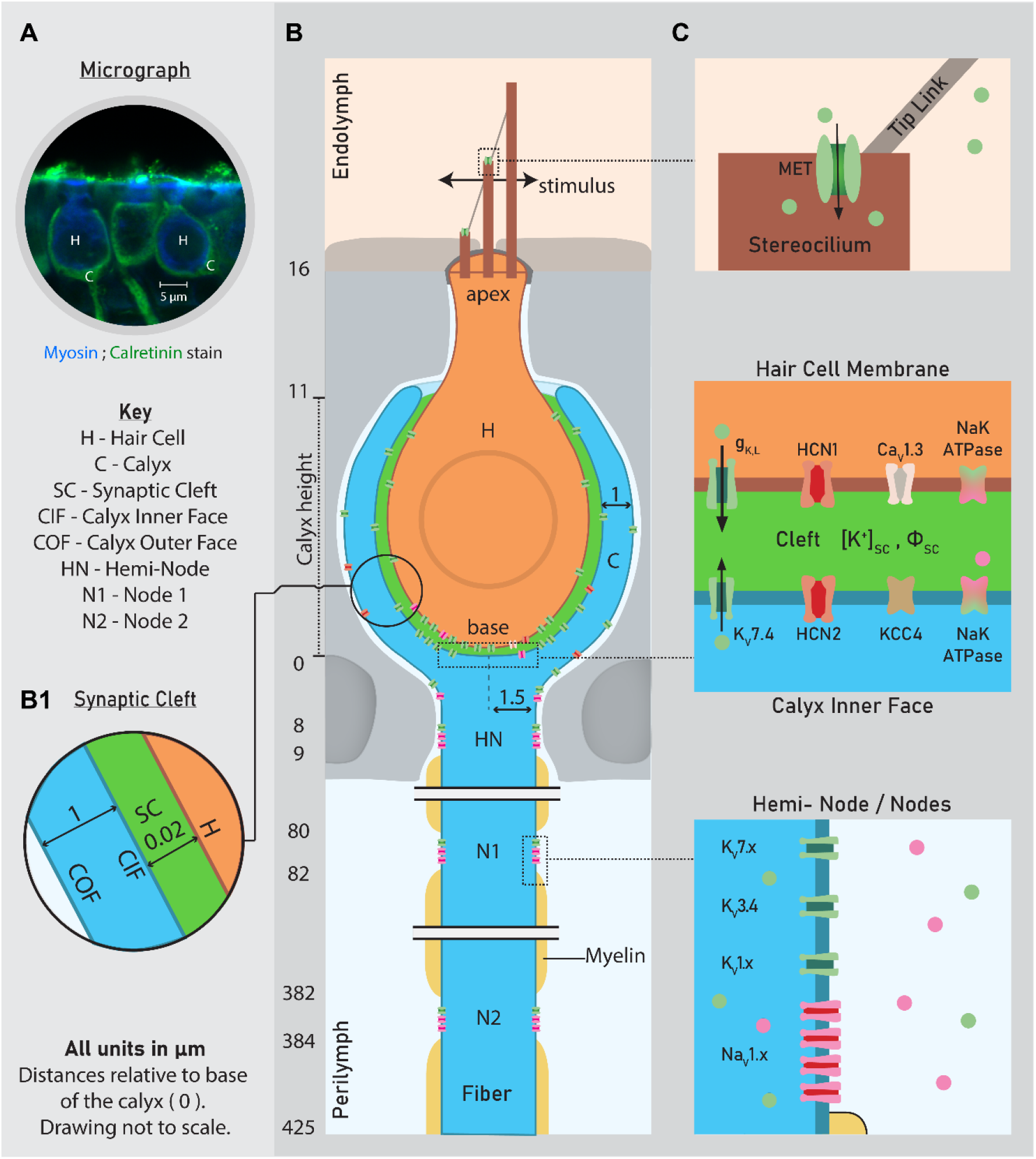
Illustration of the VHCC synapse depicting the distribution of ion channels and transporters in the hair cell (H), synaptic cleft (SC), calyx (C) and afferent fiber (F). **A,** Confocal micrograph of type I hair cells (H, blue myosin immunolabel) within calyces (C, green calretinin immunolabel). Long-Evans adult rat utricle, striolar zone. **B,** Geometry of a type I hair cell-calyx synapse from central/striolar zones of rodent vestibular organs, and the dimensions used in the model. A 2D curve representing the hair cell, synaptic cleft and calyx was generated (**SI, Fig. S1**) based on measurements from Lysakowski et al. 2011, their Fig. 2C. Long-Evans adult rat crista, central zone. Dimensions were comparable (within 10%) to those of hair cells and calyces in A. **B1,** Cleft and calyx width. **C,** The ion channel types at different key locations in the model. See **SI, Tables S6** and **S7** for sources of channel kinetics and distributions. *ϕ_SC_*, electrical potential in the synaptic cleft; [***K***^+^]_***SC***_, K^+^ concentration in the synaptic cleft.

The model incorporates ion channels and transporters identified from experimental studies of type I hair cells and associated afferents to simulate the voltage response of the calyx to mechanical and electrical stimuli: hair bundle deflection and hair cell voltage steps. Most of the parameters in the model are from published literature (**SI Tables S5, S6 and S7**); methods are provided for the unpublished data (**SI Experimental Methods**). The model was created to understand how currents and driving forces interact at the VHCC synapse during nonquantal transmission (NQT) – a change in the calyx potential in response to a change in hair cell potential that does not involve exocytosis of packets (vesicles or quanta) of neurotransmitter nor gap junctions. We do not consider quantal transmission, as the goal is to understand NQT as an independent phenomenon. In addition, mammalian calyces recorded in vitro often show only NQT. The predictions of the model are compared to experimental results in which only NQT was present at the synapse. Simulations used physiological parameters from experimental data performed at room temperature.

The key parameters of interest are changes that occur in the electrical potential and K^+^ concentration of the synaptic cleft, which is not experimentally accessible. The model incorporates capacitive and resistive currents across both the pre-synaptic hair cell and post-synaptic calyx membranes and uses continuity and electro-diffusion equations to describe the relationship between currents, electrical potentials and ion concentrations in the synaptic cleft. Using a finite element approach and COMSOL Multiphysics® software, we simulate the effects of changes in electrical potential in each model compartment and how these changes represent the mechanosensory signal as it travels from one compartment to the next. In the synaptic cleft, spatiotemporal changes in both electrical potential and ion concentrations are simulated and result in spatially varying driving forces for currents along the basal-to-apical extent of hair cell and calyx inner face membranes. Initial and boundary conditions and full details on channel kinetics are provided in **SI**. The overall geometry, localization of ion channels and other relevant variables are presented in **Fig. 1**.

## Results

We use the VHCC model to delineate biophysical events and driving forces that underlie NQT and describe their effects on the mechanosensory signal communicated to the calyx terminal of the afferent neuron. We first describe the resting state of the VHCC compartments, then responses to step bundle deflections. In each condition, we provide the model’s output in terms of: 1) electrical potentials relative to a distant ground for the hair cell (*ϕ_H_*), synaptic cleft (*ϕ_SC_*), and calyx (*ϕ_C_*); 2) trans-membrane voltages for the hair cell (*V_H_*) and the calyx inner face (*V_CIF_*) membranes facing the synaptic cleft; 3) K^+^ and Na^+^ concentrations in the synaptic cleft ([*K*^+^]_*SC*_, [*Na*^+^]_*SC*_); and 4) key ion-channel currents across each membrane. The contributions of *ϕ_SC_* and [*K*^+^]_*SC*_ are then analyzed separately to determine their roles in driving pre- and post-synaptic currents and thus NQT. We then compare model results to published data showing NQT during sinusoidal hair bundle displacement and paired voltage clamp recordings from hair cells and calyces to identify features that corroborate the existence and role of an electrical potential in the cleft. Finally, to understand the extent to which NQT depends on calyx morphology, we analyze the effect of calyx height on the electrical and potassium potentials in the cleft (*ϕ_SC_, E_K_*) and the response latency of the calyx.

### Resting state of the system

The model was fully defined with no free parameters that required additional estimation. From the initial values and boundary conditions, we calculated the resting currents, ion concentrations, electrical potentials, and membrane voltages, using the stationary solver in COMSOL (***SI, Computational Methods**).* At rest, the MET channels had a 10% open probability, corresponding to −43 pA of current. The model produced hair cell and calyx resting potentials (*ϕ_H_*: –78 mV, *ϕ_C_*: –65 mV) that are consistent with reported resting *ϕ_H_* and *ϕ_C_* (Rennie and Correia 1994; Rennie et al. 1996; Rüsch and Eatock 1996; Chen and Eatock 2000; Sadeghi et al. 2014; Spaiardi et al. 2017). Model resting output values for which there are no experimental data for comparison are: *ϕ_SC_* at the base of the synapse, +2 mV; [*K*^+^]_*SC*_ and [*Na*^+^]_*SC*_ at the base of the synapse, 7 mM and 129 mM, respectively. The non-zero value for *ϕ_SC_* emerges from currents at rest carried by the same ion channels and transporters that participate in stimulus-evoked NQT.

These results were taken as the initial conditions from which to determine, with the time-dependent solver (COMSOL), changes in response to a stimulus. In all figures, the values at t = 0 show conditions at rest.

### Synaptic response to a step displacement of the hair bundle

In **Fig. 2**, we illustrate the simulated response of sequential stages in the VHCC model to a positive (excitatory) step (+1 μm) of hair bundle displacement at 50 ms followed by a negative (inhibitory) displacement (–1 μm). The stimulus (**Fig. 2A**) is bundle displacement viewed from above relative to resting position, taken mid-way up the bundle and parallel to the plane of the epithelium (**Fig. 1B**, ‘stimulus’). The mechanotransduction current (I_MET_) elicited by bundle stimulation is shown in **Fig. 2B**. Positive bundle displacement evoked a peak inward I_MET_ of ~400 pA, and the negative step reduced I_MET_ to 0. These changes in I_MET_ depolarized and hyperpolarized *ϕ_H_* (**Fig. 2C**, modeled by **SI Eq. S2**), respectively, and altered hair cell basolateral (pre-synaptic) currents (**Fig. 3**). This model does not consider transducer adaptation, such that any decay (adaptation) of I_MET_ is driven by changes in *ϕ_H_*, which changes the driving force on I_MET_. NQT manifests as a change in calyx potential, *ϕ_C_* (**Fig. 2G**), in response to changes in hair cell potential, *ϕ_H_*. For this bundle deflection, the NQT-induced change in *ϕ_C_* was large enough to trigger an action potential (AP) in the calyx which is in turn transmitted retrogradely by NQT back to the hair cell, showing up as transients at each stage (**Fig. 2B-F, arrowheads** and **Fig. 3A-C, vertical dashed line and asterisks**).

**Fig. 2.**
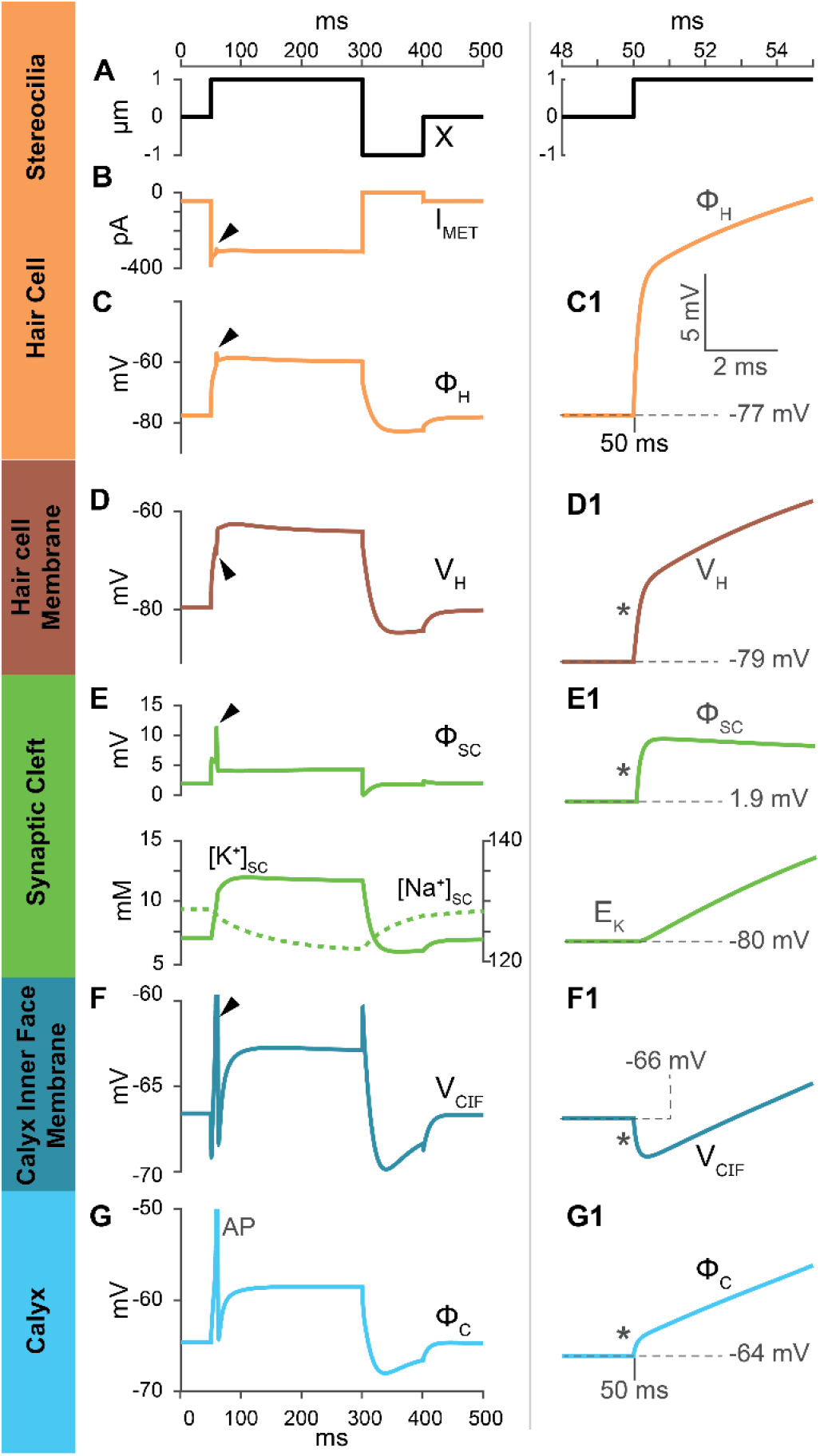
Propagation of the mechanosensory signal evoked by hair bundle displacement, illustrating dynamic changes in electrical and K^+^ potentials at each stage. **A,** Displacement (X) of the hair bundle: +1-μm, 250-ms step, beginning at t=50 ms and followed by negative step to −1-μm (300-ms) step. **B-G,** Responses to full stimulus, progressing from I_MET_ (B) to calyx postsynaptic potential (G). **Arrowheads** point to the retrograde echo of a calyx action potential that was stimulated by anterograde transmission. **C-G,** Values shown are at the base of the synapse. **C1-G1**, Responses in C-G to +step on expanded time scale (between 48 and 55 ms). **B,** I_MET_. **C, C1,** Hair cell receptor potential (electrical potential re: ground, *ϕ_H_*) rises rapidly. **D, D1,** Voltage across the basolateral hair cell membrane (*V_H_*) rises rapidly but is smaller than *ϕ_H_* by the cleft potential (*V_H_* = *ϕ_H_* – *ϕ_SC_*). **E,** [*K*^+^]_*SC*_ (*solid line;* left axis) rises while [*Na*^+^]_*SC*_ (*dotted line*; right axis) falls. **E1,** Synaptic cleft electrical potential relative to ground (*ϕ_SC_*) rises much faster than K^+^ potential (*E_K_*) for [*K*^+^]_*SC*_. **F, G,** Voltage across calyx inner face membrane (*V_CIF_* = *ϕ_C_* – *ϕ_SC_*) rapidly hyperpolarizes (**F,F1**) and then reverses as *ϕ_C_* depolarizes (**G,G1**), ultimately producing a calyceal AP (**G**). Note that after +step onset, fast components (*) of V_H_ (**D1**) and V_CIF_ (**F1**) have opposite polarities and same time course as fast rise in *ϕ_SC_* (**E1**).

**Fig. 3.**
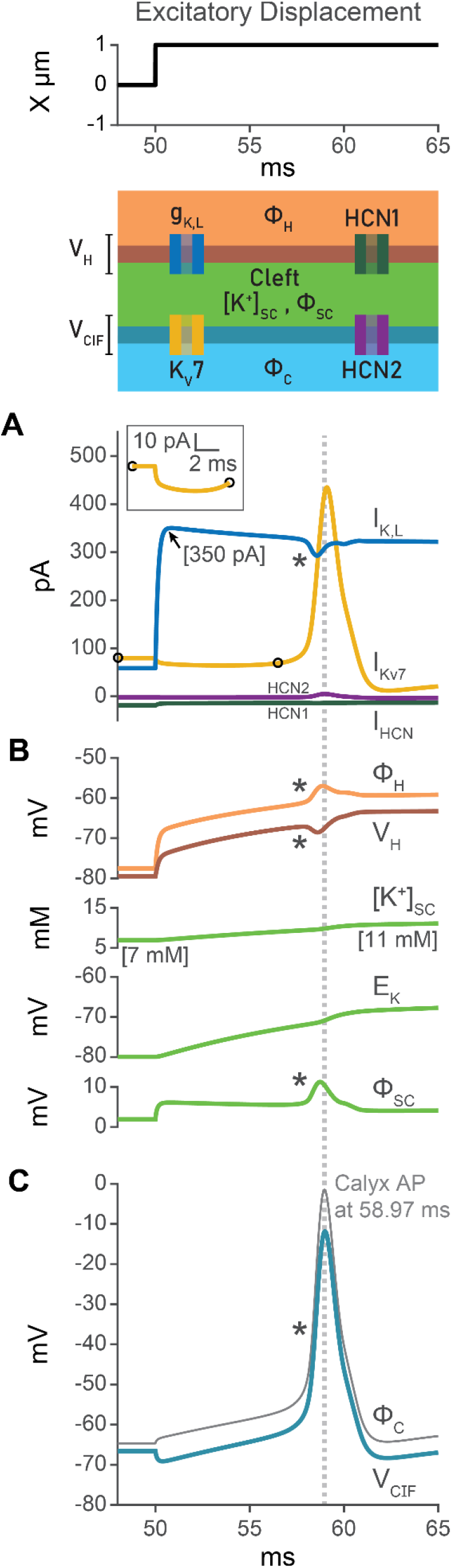
Ionic currents during hair bundle stimulation and Retrograde NQT of a calyx AP to the hair cell. During excitatory step bundle displacements (top), ionic currents (**A**), primarily through hair cell g_K,L_ and calyx K_V_7, are affected by and affect cleft properties (ϕ_SC_ and [K^+^]_SC_; **B**). Anterograde NQT evokes a single AP in the calyx (gray line, **C**; peak shown by dashed line). **Asterisks** in A-C show the retrograde influence of the calyx AP on hair cell and calyx currents (A), hair cell and cleft properties (B), and calyx (C) voltages. **A,** Net currents through g_K,L_ (hair cell, blue), K_V_7 (calyx, gold) and HCN channels (hair cell, brown, and calyx, purple) in response to the first 15 ms of the +1-μm step. **A inset,** expanded view of K_V_7 current decrease at + step onset. **B,** ϕ_H_, V_H_, [K^+^]_SC_, E_K_ and ϕ_SC_ at the base of the synapse during excitatory stimulus, showing that fast retrograde transmission to the hair cell occurs through ϕ_SC_ and not E_K_. See **asterisks**. **C,** V_CIF_ (cyan) and ϕ_C_ (gray, featuring the calyx AP) at the base of the synapse during excitatory stimuli.

### Predicted Changes in Cleft Electrical Potential, Na^+^ and K^+^ Concentrations and Transmembrane Voltages

The model permits simulation of biophysical properties of the cleft microenvironment, which is not experimentally accessible. During NQT, the synaptic cleft potential, *ϕ_SC_*, changes as shown in **Fig. 2E,E1.** The dynamic changes in *ϕ_SC_* (modeled by **SI Eq. S5**) allow calculation of pre-synaptic (*V_H_* = *ϕ_H_* – *ϕ_SC_*) and post-synaptic (*V_CIF_* = *ϕ_C_* – *ϕ_SC_*) trans-membrane voltages. *V_H_* (**Fig. 2D**) and *V_CIF_* (**Fig. 2F**) differ by *ϕ_SC_* from the intracellular potentials with respect to ground (*ϕ_H_* and *ϕ_C_*, **Fig. 2C,G**).

**Figure 2E** shows the predicted changes in [*K*^+^]_*SC*_ and [*Na*^+^]_*SC*_ in the synaptic cleft at its base: K^+^ accumulation was accompanied by Na^+^ depletion, as predicted by a previous electrodiffusion model of the type I-calyx that considered the effect of the cleft on the hair cell receptor potential (Soto et al. 2002). In our VHCC model, changes in ion concentrations in the cleft are caused by: 1) altered ionic currents, mostly through channels in the pre-synaptic and post-synaptic membranes; and 2) the axial electro-diffusion of ions from the source (ion channels in the depth of the cleft) to the sink (apical edges of the calyx) (**Fig. 1**, and **SI Eqs. S3,S4**).

#### Synaptic cleft electrical potential, *ϕ_SC_*, rises much faster than the synaptic K^+^ potential

To more clearly illustrate the variables responsible for the speed of NQT we show the dynamic changes in *ϕ_H_* (**Fig. 2C1**) and *ϕ_C_* (**Fig. 2G1**), *V_H_* and *V_CIF_* (**Fig. 2D1,F1**), and synaptic cleft potential, *ϕ_SC_* (**Fig. 2E1**) at the onset of the +1 μm step. We also plot the equilibrium potential of K^+^, (*E_K_*), which reflects changes in synaptic cleft potassium level, [*K*^+^]_*SC*_ (**Fig. 2E1**).

Differences in the time courses of stimulus-evoked changes in *ϕ_SC_*, *E_K_*, and *V_CIF_* provide key insight into the driving forces for NQT. At the onset of the positive step, the synaptic cleft potential, *ϕ_SC_*, rises from 1.9 mV to +6 mV in 0.43 ms, 8-fold faster than the time for *E_K_* to rise the same amount (−79.9 mV to −75.8 mV) (**Fig. 2E1**). The rise in *E_K_* is also slow compared to the change in *V_CIF_* (**Fig. 2F1**). This provides an important insight into the mechanism of NQT that has not been previously recognized: *ϕ_SC_* provides the initial driving force for change in the post-synaptic membrane current, and only later does change in *E_K_* make a significant contribution. This difference arises because fewer ions are needed to change *ϕ_SC_* than *E_K_*. **Fig. 2C1-F1** highlight how changes in *ϕ_SC_* (**Fig. 2E1**) affect the magnitude of *V_H_* (**Fig. 2D**) and the magnitude and direction of *V_CIF_* (**Fig. 2F**). At step onset, *V_CIF_* hyperpolarizes while *ϕ_C_* depolarizes (**Fig. 2F1,G1, asterisks)**. The hyperpolarization of *V_CIF_* occurs because the fast depolarization of the cleft (*ϕ_SC_*) is larger than the fast depolarization of the intracellular calyx potential (*ϕ_C_*) making *V_CIF_* = (*ϕ_C_* – *ϕ_SC_*) negative. The subsequent slow rise in *V_CIF_* follows the depolarization of *ϕ_C_* which results from the slow increase in [*K*^+^]_*SC*_ and *E_K_*. That *V_CIF_* and *ϕ_C_* are different and can change in opposing directions makes it apparent that voltage across the calyx inner and outer face membranes can be different despite the common intracellular potential. While the VHCC model results agree with experiments showing that changes in synaptic cleft *[K^+^]* affect both hair cell and calyx signals (Lim et al. 2011; Contini et al. 2012, 2017; Spaiardi et al. 2020), the model further shows a pivotal role for the synaptic cleft potential in reducing transmission latency. The independent contributions of *ϕ_SC_* and [*K*^+^]_*SC*_ to NQT are analyzed later.

### Contributions of Different Ion Channels to Nonquantal Transmission

The synaptic cleft is a dynamic system where electric potentials, ion concentrations and ionic currents interact. The changes in cleft electrical potential and ion concentrations shown in **Fig. 2** are driven by currents through voltage-sensitive ion channels (see **SI, Table S6** for kinetics) on the hair cell basolateral membrane and on the calyx inner face, and in turn modulate these currents (**Fig. 3**). NQT is bidirectional: we first describe the roles of key channels during anterograde (hair cell to calyx) NQT and later discuss retrograde (calyx to hair cell) NQT.

#### Currents through open g_K,L_ channels respond very rapidly to hair cell depolarization

The large low-voltage-activated K^+^ conductance “g_K,L_” is a distinctive feature of type I vestibular hair cells (Correia and Lang 1990; Rennie and Correia 1994; Rüsch and Eatock 1996). At the resting bundle position (t<50 ms, **Figs. 2** and **3**) before the step, *V_H_* is only slightly positive (~1 mV) of *E_K_* (−80 mV; **Fig. 2E1**). Despite this small driving force, g_K,L_ has a large open probability (~50%) because of its very negative activation range and carries significant outward K^+^ current, I_K,L_, into the cleft (**Fig. 3A**). At the onset of the positive step, *ϕ_H_* and *V_H_* depolarize faster than *E_K_* (**Fig. 2C1-E1**). Since g_K,L_ is already significantly open, the increase in driving force (*V_H_* – *E_K_*) instantaneously increases I_K,L_ (**Fig. 3A**), altering both *ϕ_SC_* and [*K*^+^]_*SC*_. I_K,L_ is largely sustained; a modest decrease (~10%, from +350 pA to +317 pA) reflects the reduction in driving force as [*K*^+^]_*SC*_ increases and *E_K_* depolarizes (**Fig. 3B**).

In **Fig. 3** we have presented local potentials at the base of the synapse and whole-cell ionic currents. However, the VHCC model has the ability to calculate the spatio-temporal distributions of these variables. In our model conditions, the net I_K,L_ was never inward (negative). However, by using the model to examine membrane currents as functions of location along the cleft, we noticed that inward I_K,L_ can occur locally at the base of the hair cell during a negative step if *V_H_* becomes negative to *E_K_* (**SI Fig. S2**). These results support the idea that g_K,L_ may under certain conditions carry inward K^+^ currents into the hair cell and aid in K^+^ clearance (Spaiardi et al. 2020).

**Figure 3** shows that the current through g_K,L_ is fast, large, and non-inactivating, so that it closely follows the hair bundle stimulus. Since g_K,L_ is high in the resting state, the receptor potential-induced change in I_K,L_ is not delayed by ion channel activation and is near instantaneous. I_K,L_ rapidly changes the cleft electrical potential (**Fig. 3B,** *bottom trace*), which in turn rapidly alters currents through low-voltage-activated potassium (K_LV_) channels in the calyx inner face membrane.

#### Excitatory stimuli decrease K_V_7 current into the synaptic cleft from the calyx

The postsynaptic K_LV_ conductances include the K_V_7 family of voltage-gated channels (Hurley et al. 2006). The VHCC model permits quantitative characterization of the role of these channels in modulating [*K*^+^]_*SC*_ and *ϕ_SC_* during NQT, which has not previously been clear. According to the model, at resting membrane potential (*V_CIF_* = −68 mV), outward current through partly open (29%) K_V_7 channels on the calyx inner face supplies K^+^ to the synaptic cleft, helping to set resting levels of [*K*^+^]_*SC*_ and *ϕ_SC_*. During positive bundle displacement, changes in *V_CIF_* and *E_K_* across the calyx inner face membrane (**Fig. 3B**) slightly reduce outward K_V_7 currents into the synaptic cleft (**Fig. 3A,** I_KV7_) except during the calyx action potential (AP) where outward K_V_7 currents rise (**Fig. 3A,** I_KV7_, asterisk).

#### Synaptic HCN channels establish resting conditions but have minimal effect on [*K*^+^]_*SC*_ and *ϕ_SC_* during NQT

Hyperpolarization-activated cyclic-nucleotide-gated (HCN) channels are found on both sides of the VHCC synapse. Type I hair cells primarily express HCN1 channel subunits (Horwitz et al. 2011) and vestibular afferent neurons primarily express HCN2 subunits (Horwitz et al. 2014). Model results show that under physiological concentrations and potentials, HCN channels support a net inward current in both cells, mediated by influx of Na^+^ ions, that is 10-100-fold smaller than outward K^+^ currents through g_K,L_ and K_V_7 channels (**Fig. 3A**). In the model, we localized HCN1 to the hair cell and HCN2 to the calyx and found that both HCN channels had smaller open probabilities (HCN1-18%; HCN2-8%) than K_LV_ channels (g_K,L_ - 50%; K_V_7 - 29%) at rest (*V_H_* = −79 mV; *V_CIF_* = −68 mV) and contributed very small currents to either positive (**Fig. 3A**) or negative (**SI Fig. S2**) step responses. This difference relative to K_LV_-associated currents is explained by the HCN channels’ smaller whole-cell conductance values (**see SI, Table S6**), their hyperpolarized voltage range of activation, and their mixed cationic currents (K^+^ and Na^+^) moving in opposite directions. In the model, both HCN1 and HCN2 channels, along with the Na^+^/K^+^-ATPase, contribute to the regulation of [K^+^] and [Na^+^] in the synaptic cleft (**Fig. 2E**). Blocking HCN channels has been experimentally shown to block NQT (Contini et al. 2017), but only under conditions that artificially favor HCN channels: i.e., when the calyx was held at −100 mV, an unphysiological (substantially negative to *E_K_*) condition at which most K_V_7 channels would be closed and most HCN channels would be open.

#### Retrograde transmission from calyx to hair cell

The VHCC model naturally captures retrograde transmission (**Fig. 2B-F, arrowheads**), defined as a change in hair cell potential caused by a change in calyx potential. This is especially pronounced during NQT-induced postsynaptic action potentials that occur for sufficiently large hair bundle displacements. An action potential in the fiber (**Fig. 2G; Fig. 3C**) causes the calyx inner face membrane to depolarize (**Fig. 3B,** V_CIF_) and increases outward current, primarily through K_V_7 channels (**Fig. 3A**; compare K_V_7 and HCN2); *ϕ_SC_* rises (**Fig. 3B**); *V_H_* becomes more negative (**Fig. 3B**); outward K^+^ currents from the hair cell, primarily through g_K,L_ channels, decrease (**Fig. 3A,** see g_K,L_ and HCN1); and the hair cell potential, *ϕ_H_,* depolarizes (**Fig. 2C, arrowhead; Fig 3B**). The fast component of retrograde transmission is mediated by changes in *ϕ_SC_* and not *E_K_* (**Fig. 3, asterisks**). These results indicate that fast retrograde events seen in electrophysiological recordings of the hair cell and calyx (Contini et al. 2017) are caused by changes in electrical potential in the synaptic cleft. It has been suggested that the bidirectional nature of nonquantal transmission, which our VHCC model captures, could be used to modulate the sensitivity of both the calyx and the hair cell (Holt et al. 2017).

### Role of the Synaptic Cleft in Nonquantal Transmission: *ϕ_SC_* and [*K*^+^]_*SC*_

The predictions of the VHCC model agree with experimental observations (Lim et al. 2011; Contini et al. 2012, 2017, 2020; Spaiardi et al. 2020) that support the hypothesis that K^+^ currents through the basolateral membrane of the type I hair cell modulate [*K*^+^]_*SC*_ (Chen 1995; Goldberg 1996) and change *E_K_* to affect pre- and post-synaptic currents. The model further shows that modulation of *ϕ_SC_* also alters both pre-synaptic and post-synaptic ionic currents but on a substantially faster time scale. What are the relative contributions of *ϕ_SC_* and [*K*^+^]_*SC*_ to NQT? Are both mechanisms necessary? What advantages do they provide? To answer these questions, we use the VHCC model to isolate *ϕ_SC_* and [*K*^+^]_*SC*_ during both step and sinusoidal bundle displacements.

#### Isolation of the contributions of *ϕ_SC_* and [*K*^+^]_*SC*_ during step bundle displacements

We mathematically define two conditions: *ϕ_SC_* modulation and [*K*^+^]_*SC*_ modulation. The *ϕ_SC_* modulation-only condition is set by fixing [*K*^+^]_*SC*_ and [*Na*^+^]_*SC*_ at 5 mM and 140 mM, respectively, while *ϕ_SC_* is allowed to vary as a function of ionic currents (**SI Eqs. S8-10**). The [*K*^+^]_*SC*_ modulation-only condition is set by fixing *ϕ_SC_* at 0 mV and allowing [*K*^+^]_*SC*_ and [*Na*^+^]_*SC*_ to vary as functions of the ionic currents (**SI Eqs. S11-13**). **Figure 4** illustrates how *ϕ_SC_* and [*K*^+^]_*SC*_ separately and together shape the calyx potential (**Fig. 4B1,B2**) and its time derivative (**Fig. 4C1,C2**) at the onset of 2 positive bundle displacements. The *ϕ_SC_* modulation reduced the latency of the calyx potential by 1.8 ms (0.3 μm step) and 1.5 ms (1 μm step). **Figures 4C1,C2** show the considerably greater rate of change in calyx potential, 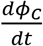, for the *ϕ_SC_* condition compared to the [*K*^+^]_*SC*_ condition, and how both combine.

**Fig. 4.**
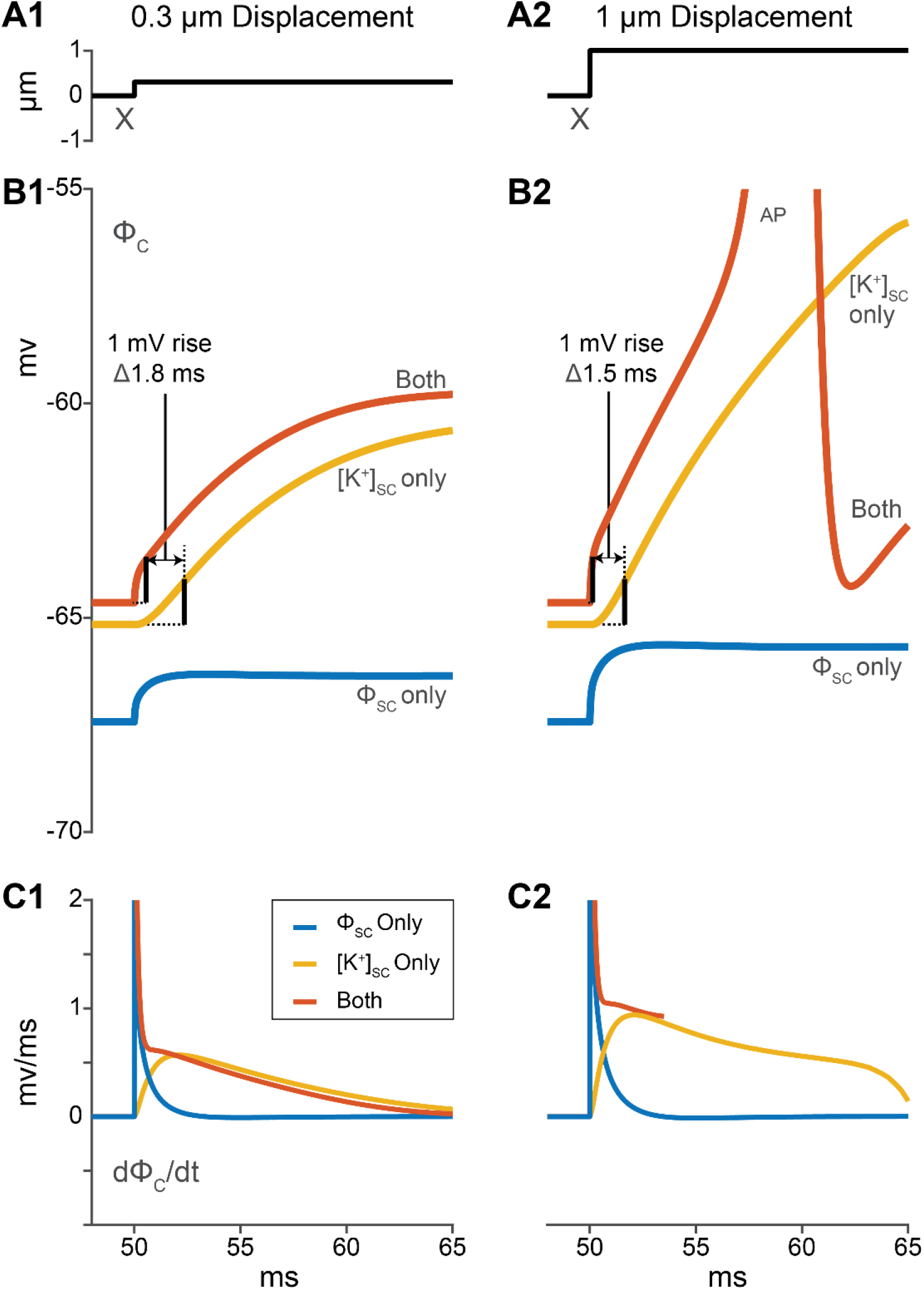
*ϕ_SC_* and [*K*^+^]_*SC*_ are responsible for the fast and slow components of NQT respectively. Simulations for 2 bundle steps (**A1, A2**) with *ϕ_SC_* modulation only (blue), with [*K*^+^]_*SC*_ modulation only (yellow) and with both enabled (dark orange). **B1,B2,** Calyx potential (*ϕ_C_*) rises faster when *ϕ_SC_* is present, either alone or together with [*K*^+^]_*SC*_. The time to rise by 1 mV (dashed lines) clearly shows the lag in K^+^ accumulation. Together, ‘Both’ changes trigger a calyx AP for the larger stimulus. **C1,C2,** Rate of change of calyx potential highlights the kinetic differences in *ϕ_SC_* and [*K*^+^]_*SC*_ changes. For clarity, the ‘Both’ curve (red) is truncated before the AP in **C2**.

**Figure 4** also shows that [*K*^+^]_*SC*_ modulation is responsible for the slow component of the rise in calyx potential. When *ϕ_SC_* and [*K*^+^]_*SC*_ are allowed to vary concomitantly, both fast and slow components are seen. Comparing the rise of *ϕ_C_* in the [*K*^+^]_*SC*_-only and ‘Both’ conditions clearly shows that change in *ϕ_SC_* confers a ~1-2 ms advance to the calyx response (**Fig. 4B**). Thus, the modulation of *ϕ_SC_* enables faster signal transmission between hair cell and calyx. The modulation of [*K*^+^]_*SC*_, although slower, results in a larger depolarization of the calyx. For these inputs to the VHCC model, both mechanisms together did not trigger spiking for smaller hair bundle deflections, but both were necessary to elicit spiking for the larger deflection. In the presence of an action potential, the response of the model at each stage is influenced by currents at the hemi-nodes and nodes of the afferent fiber (**SI, Eqs. S6, S7** and **Table S6**). *ϕ_C_* (**Fig. 4B2**) closely resembles experimental whole-cell recordings of the voltage response of a rat utricular calyx to positive step displacements (Songer and Eatock 2013 – their Fig. 5A).

**Fig. 5.**
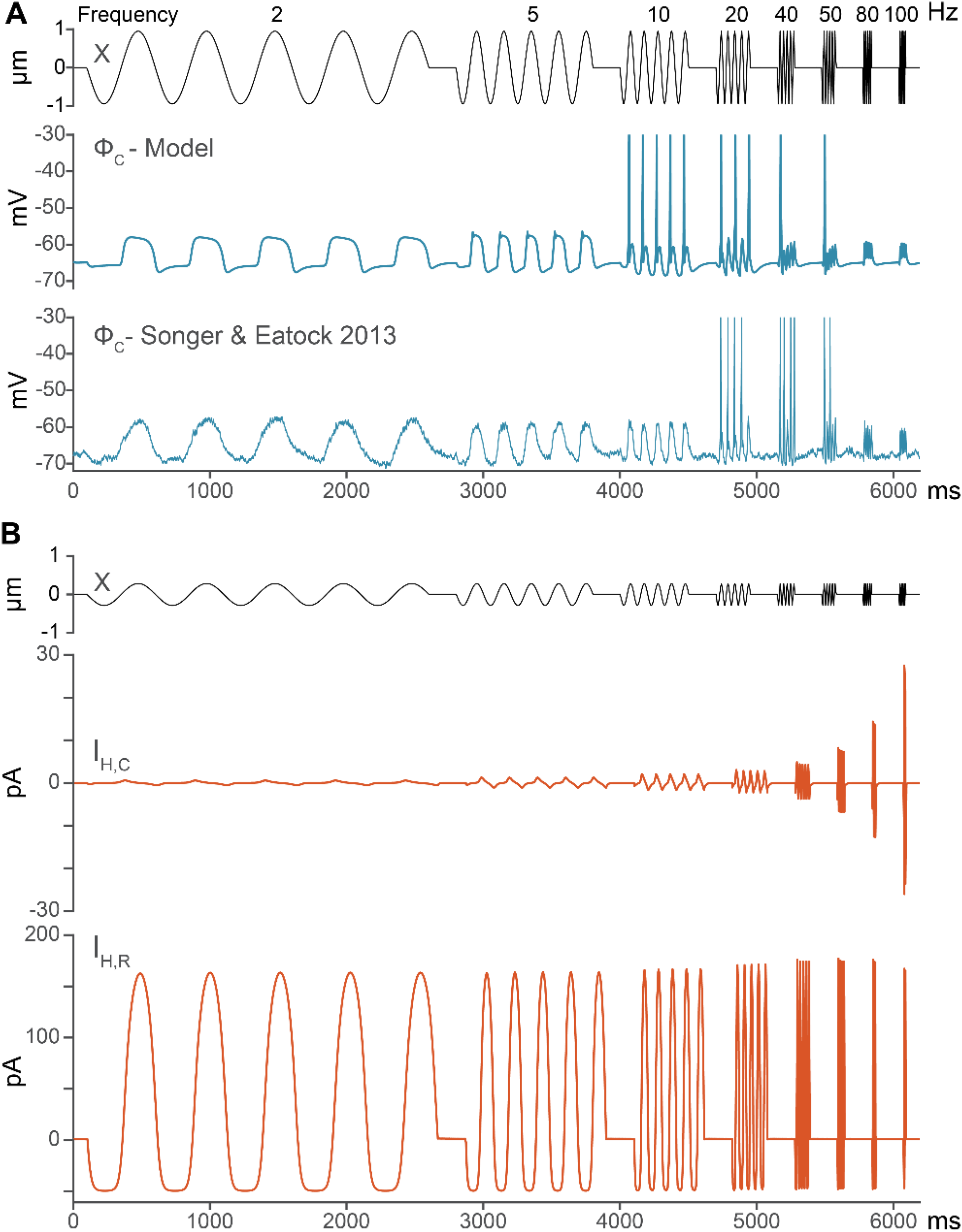
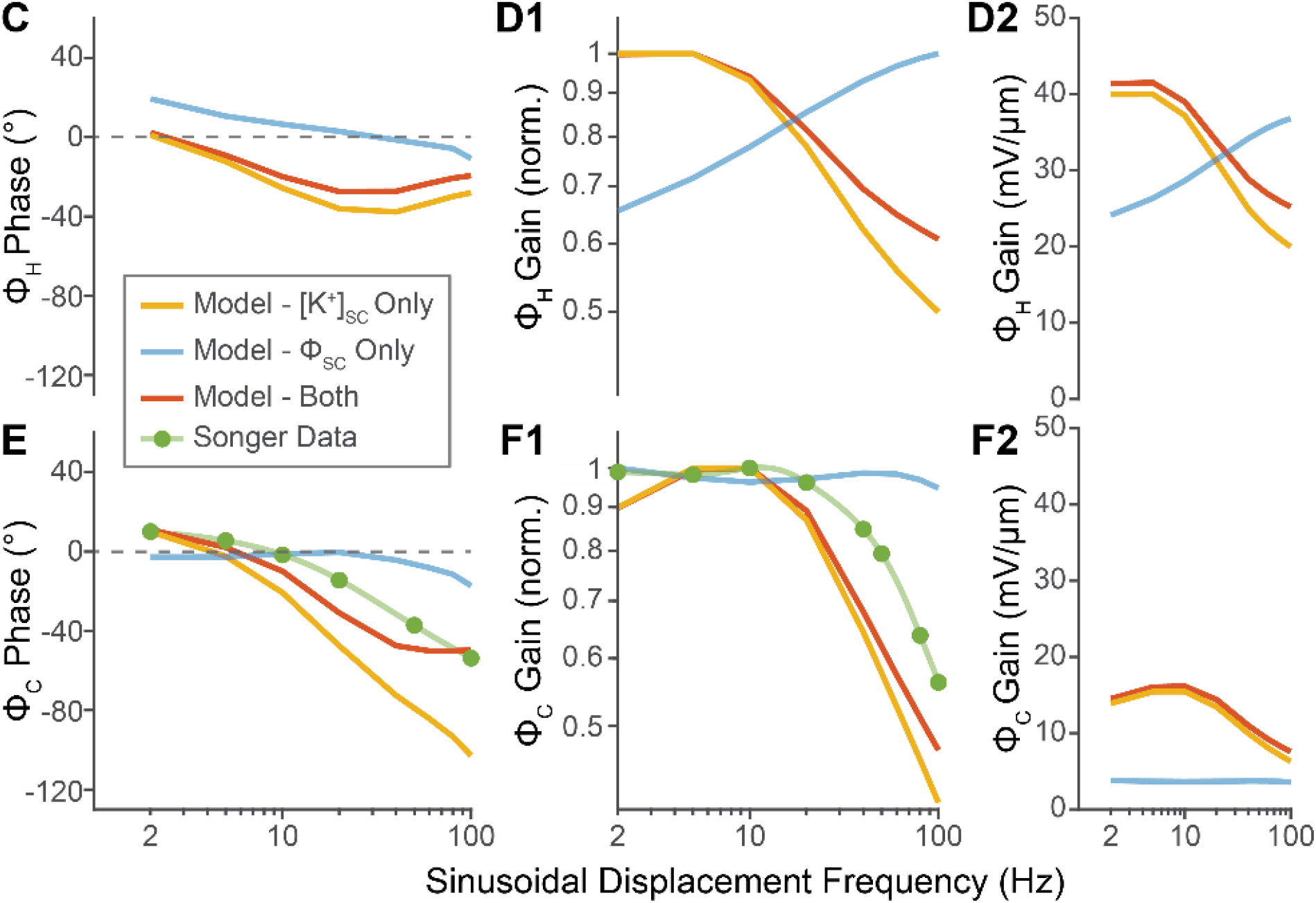
Resistive currents from the hair cell drive changes in *ϕ_SC_* and [*K*^+^]_*SC*_ to shape the phase and gain of the transmitted signal (*ϕ_C_*). **A,** Modelled *ϕ_C_* response and whole-cell current clamp recording of *ϕ_C_* from rat saccular calyx (Songer and Eatock 2013, their Fig. 5B) during ±1 μm sinusoidal bundle displacement. **B-F,** Model outputs for ±0.3 μm bundle displacement analyzed into *ϕ_SC_* modulation-only component (*blue*), [*K*^+^]_*SC*_ modulation-only component (*yellow*) and *Both* together (*dark orange*). **B,** Capacitive (I_H,C_) and resistive (I_H,R_) currents from the hair cell. I_H,R_ is the primary contributor to changes in the synaptic cleft. **C-F,** Bode plots showing phase (**C, E)**, normalized gain (**D1, F1)** and gain (**D2, F2**) of *ϕ_H_* and *ϕ_C_* relative to X. **E,** [*K*^+^]_*SC*_ component (yellow) falls off much more steeply and with lower corner frequency than *ϕ_SC_* component (blue). Combining them (dark orange) better matches experimental data (green, from Songer and Eatock 2013, their Fig. 5C). **F1,F2,** [*K*^+^]_*SC*_ modulation dominates the gain of NQT between 2-100 Hz. Allowing *ϕ_SC_* to also change (dark orange) slightly increases gain at higher frequencies (yellow, **F1**). *ϕ_SC_* provides a small constant gain across a wide frequency band (**F2**).

### The VHCC Model Captures the Frequency Response and Gain of NQT

Songer and Eatock (2013) recorded responses from calyces in the excised rat saccule to sinusoidal hair bundle stimulation between 2-100 Hz in order to capture the frequency filtering characteristics relevant to natural head motions (Cullen 2019). An example of NQT from these experiments is reproduced in **Fig. 5A.** In addition to measuring the frequency-following capacity of the NQT-induced changes in calyx potential (*ϕ_C_*), these data showed that for large bundle displacements NQT alone (without quantal transmission) can trigger action potentials in the afferent. Here we show the VHCC model’s ability to reproduce these salient features of the experimental data and use the model to illustrate the roles of [*K*^+^]_*SC*_ and *ϕ_SC_* in shaping the calyx response (**Fig. 5**).

#### The model captures NQT-driven firing in response to sinusoidal stimulus

The model produced calyx potentials that are consistent with whole-cell recordings from a calyx in the central striolar zone of the rat saccular epithelium (**Fig. 5A**), including spikes for large stimuli. In both data and model, the calyx was most likely to spike for stimulus frequencies between 10 and 50 Hz.

#### Resistive currents dominate capacitive currents during bundle stimulation

The VHCC model enables us to track the contributions of capacitive and resistive currents to NQT as a function of frequency (**SI Eqs. S3-5, S8-13**). In **Fig. 5B** and **5C** a small stimulus (±0.3 μm) was used to avoid triggering APs and better reveal the underlying changes in hair cell current and postsynaptic potentials (*ϕ_C_*). The model predicts resistive currents (I_H,R_) are much larger than capacitive current (I_H,C_) flowing from the type I hair cell (**Fig. 5B**, note the different y-axis limits). The capacitive current does increase with frequency, as expected, and can noticeably affect NQT during large voltage step stimuli (discussed later in Figure 6).

**Fig. 6.**
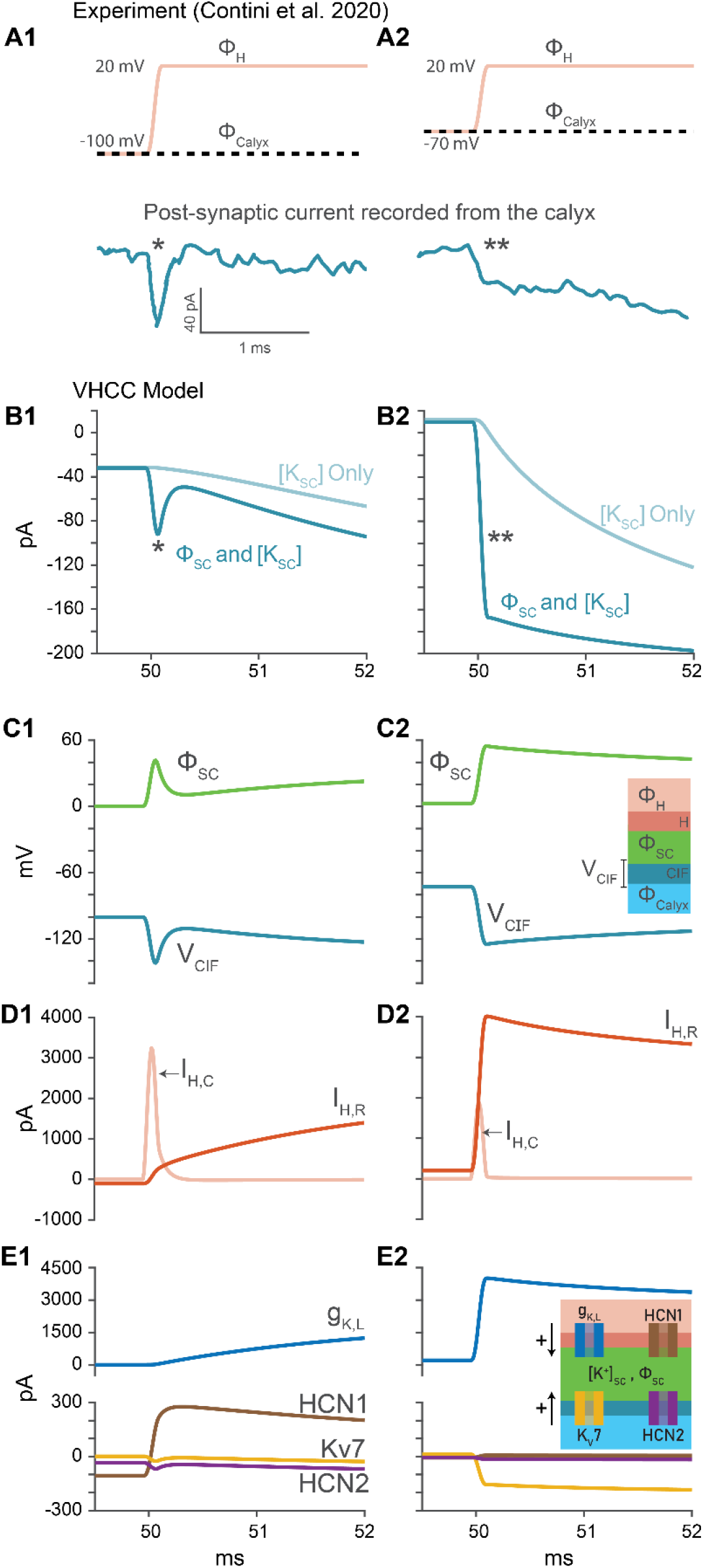
Changes in synaptic cleft potential *ϕ_SC_* explain the shape of fast post-synaptic currents in paired voltage recordings. **A1,A2,** Voltage protocols and currents from experiment. Calyx Potential (*ϕ_C_*) held at −100 mV. Hair cell potential stepped at 50 ms from holding potential of −100 mV to 20 mV (**A1**). Calyx Potential (*ϕ_Calyx_*) held at −70 mV. Hair cell potential stepped at 50 ms from holding potential of −70 mV to 20 mV (**A2**). Currents reproduced from Contini et al. (2020, their figure 10). **B1,B2,** Resistive post-synaptic current through the calyx inner face membrane, I_CIF,R_. Note that fast currents (* and **) do not occur in the [*K*]_*SC*_-only case (faded blue). **C,** Electrical potential in the synaptic cleft (*ϕ_SC_*) and voltage across the calyx inner face (V_CIF_) membrane. **C1,** The cleft potential is primarily shaped by capacitive currents from the hair cell (I_H,C_, **D1**). **C2,** The cleft potential is primarily shaped by resistive currents from the hair cell (I_H,R_, **D2**). **E,** Whole cell currents through individual ion conductances. From the non-physio-logical −100 mV holding potential (A1), HCN1 hair-cell current is prominent, fast and sustained, because HCN channels are open at −100 mV (**E1**). Note that from −70 mV holding potential (A2), g_K,L_ current is fast and sustained and K_V_7 current (yellow) is the most prominent calyx current (**E2**). Currents through g_K,L_ (E2) sustain the step in *ϕ_SC_* following the voltage step in the hair cell. The slow decay reflects the change in driving force as K^+^ accumulates in the cleft.

#### Changes in both *ϕ_SC_* and [*K*^+^]_*SC*_ are required to explain the phase of the *ϕ_C_* response to sinusoidal stimuli

We examined the contribution of *ϕ_SC_* and [*K*^+^]_*SC*_ to the phase of *ϕ_H_* (**Fig. 5C**) and *ϕ_C_* (**Fig. 5E**) as a function of frequency. A positive phase indicates that the maximal response of *ϕ_H_* or *ϕ_C_* is achieved before the maximal bundle displacement and is indicative of a high-pass filtering process. As shown, *ϕ_SC_* contributes to a positive phase response in *ϕ_H_* at frequencies below 32 Hz. For *ϕ_H_,* at lower frequencies, the positive phase arises from g_K,L_ activation (for kinetics, see **SI Table S6**): during the depolarizing half cycle, an appreciable increase in g_K,L_ (>10% at 2 Hz) produces an increasing outward current that truncates the rise in *ϕ_H_* before the peak of the bundle displacement. Likewise, for *ϕ_C_* the positive phase at lower frequencies arises from activation of K_LV_ channels within the calyx and fiber. The *ϕ_C_* response was nearly flat with frequency for the *ϕ_SC_*-only condition but showed strong low-pass filtering for the [*K*^+^]_*SC*_-only component.

The low-pass behavior of [K^+^] modulation can be explained by the time course of K^+^ accumulation or depletion in the cleft (see **SI Eq. S3**). The influence of this effect is relatively slow because more ions are needed to change ion concentrations and chemical potentials than to change electrical potential. In contrast, modulation of *ϕ_SC_* instantaneously affects voltage across both pre- and post-synaptic membranes, altering both capacitive currents and resistive (ionic) currents through open channels. This explains the short response latency at the onset of a step stimulus (**Fig. 4B**) and the *ϕ_C_* phase response to sinusoidal stimuli. The *ϕ_SC_* component is required to closely approximate the phase of experimental data above 5 Hz (**Fig. 5E**). This effect supports the evidence from step data (**Fig. 4**) that a stimulus-modulated electrical potential in the synaptic cleft speeds up NQT.

#### [*K*^+^]_*SC*_ dominates the gain of NQT but *ϕ_SC_* slightly boosts gain at all frequencies

Next, we examined the normalized and actual gain of averaged peak *ϕ_H_* (**Fig. D1,D2**) and *ϕ_C_* (**Fig. F1,F2**) as a function of frequency. For *ϕ_H_,* the [*K*^+^]_*SC*_ mechanism contributes to a positive gain that rolls off above ~20 Hz, while the *ϕ_SC_* mechanism linearly increases with frequency. Interestingly, the *ϕ_SC_* mechanism kicks in to noticeably boost the *ϕ_H_* gain above ~30 Hz. For *ϕ_C_*, the gain at low frequencies is larger for the [*K*^+^]_*SC*_ component than for the *ϕ_SC_* component (**Fig. 5F2**). The model also shows that the [*K*^+^] modulation mechanism amplifies the calyx potential response at all frequencies but especially below 20 Hz. This effect is consistent with the increasing size of the [*K*^+^]_*SC*_ component relative to the *ϕ_SC_* component with time during a displacement step (**Fig. 4B1,B2)**. The *ϕ_SC_* component is approximately constant with frequency, and the model predicts will boost the high frequency gain (**Fig. 5F1).** However, the experimental *ϕ_C_* gain (green) has a higher corner frequency (66 Hz) than the model output (36 Hz) which may reflect differences in afferent fiber properties in the model and specific experiments.

### Paired Voltage Clamp Recordings from Hair Cell and Calyx Provide Evidence of Changes in Synaptic Cleft Potential

We have previously speculated that NQT involves the flow of K^+^ through a large number of g_K,L_ channels that are open at typical hair cell resting potentials (Eatock 2018). In this condition, receptor potentials instantaneously change K^+^ current flowing into the synaptic cleft, bypassing the time—on the order of 1-100 ms—required to activate voltage-gated channels from the closed conformation. Experiments by Contini et al. (2020), involving paired voltage-clamp recordings of whole-cell currents from turtle type I hair cells and postsynaptic calyces, address this hypothesis. In these experiments, the hair cell was stepped to +20 mV from either −100 mV or −70 mV, while the calyx was held at either −100 mV or −70 mV, and fast inward calyx currents were recorded (**Fig. 6A1,A2**, respectively). Contini et al. (2020) attribute these currents to ‘resistive coupling’ through pre-synaptic and post-synaptic ion channels open at the holding potential immediately prior to the voltage step. However, the driving forces underlying the recorded currents were unclear. Additionally, despite similar time courses and temporal proximity to the voltage step, one of the fast post-synaptic currents (**Fig. 6A1**) was thought to be an artefact. Here we show with the VHCC model that the fast post-synaptic currents at the two different holding potentials are both driven by changes in *ϕ_SC_*, the electrical potential in the synaptic cleft.

The VHCC model was applied to analyze the experimental data with either the [*K*^+^]_*SC*_-only condition or the [*K*^+^]_*SC*_ and *ϕ_SC_* (‘Both’) condition, described earlier. The fast post-synaptic currents only occurred in the ‘Both’ condition (**Fig. 6B1,B2**), illustrating that changes in *ϕ_SC_* were the primary driving force and that the currents cannot be explained solely by changes in *E_K_*. Under the ‘Both’ condition, we first examined the model’s output for a calyx and hair cell holding potential of −100 mV (**Fig. 6, Left Column**), to compare with results from Contini et al. (2017, 2020). At this holding potential, g_K,L_ and K_V_7 channels were mostly closed and carried very little current prior to the voltage step. As *ϕ_H_* is depolarized to +20 mV, *ϕ_SC_* (**Fig. 6C1**) is primarily shaped by capacitive currents across the basolateral hair cell membrane (I_H,C_) which precede g_K,L_ activation (**Fig. 6D1,E1**). This variation in *ϕ_SC_* alters *V_CIF_*, shapes currents through the calyx inner face membrane (**Fig. 6B1, asterisk**), mostly via HCN2 channels (**Fig. 6E1**), and explains the current thought to be an artefact by Contini et al. 2020 (**Fig. 6A1, asterisk**). We identify these currents as evidence for changes in *ϕ_SC_*, in response to depolarization of the hair cell.

We then examined the model’s output for a calyx and hair cell holding potential −70 mV (**Fig. 6, Right Column**). At this holding potential, g_K,L_ and K_V_7 channels were significantly open prior to the voltage step. As *ϕ_H_* is depolarized to +20 mV, *ϕ_SC_* (**Fig. 6C2**) is primarily shaped by resistive currents through the basolateral hair cell membrane (I_H,R_), mostly via g_K,L_ (**Fig. 6D2,E2**). This hyperpolarizes *V_CIF_* (**Fig. 6B2**) and increases the inward driving force for currents through the calyx inner face, principally through KV7 channels rather than HCN channels (**Fig. 6E2**). This results in a distinctive and fast current (double asterisks) of similar time course (albeit increased amplitude) to that seen by Contini et al. (2020) (compare **Fig. 6A2** and **6B2**). The difference in amplitude may reflect differences in key channel properties (e.g., the voltage dependence and size of g_K,L_) and calyx geometry between the model parameters (based on data from mice and rats) and this particular experimental data (from turtles).

Under both experimental conditions (**Fig. 6A1,A2**) it is apparent that the recorded currents have a fast (asterisk and double asterisks) and a slow component. These components are consistent with the existence of two driving forces with different time courses – *ϕ_SC_* and *E_K_*. We note that *ϕ_SC_* (**Fig. 6C1,C2**) follows the general shape of 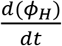 when capacitive currents dominate (at −100 mV holding condition) or *ϕ_H_* when resistive currents from the hair cell dominate, which is the case near the holding potential of −70 mV (**Fig. 6D2, Fig. 5D**) and closer to reported hair cell resting potentials (~-80 to −60 mV, see **SI Table S4** for references). Comparison of the two different experimental conditions also demonstrates that open g_K,L_ channels (**Fig. 6E2**) are responsible for the rapid and sustained transmission of the hair cell receptor potential to the calyx through changes in cleft potential and post-synaptic currents (**Fig. 6B2,C2**). These results show the importance for NQT of the unusual properties of g_K,L_: together the large magnitude and the unusually negative half-activation voltage ensure that many channels are open near the hair cell resting potential and poised to respond to incoming stimuli.

### Nonquantal Transmission Depends on Calyx Morphology

The simulations we have presented are for a calyx shape typical of many calyces in the central and striolar zones of rodent vestibular epithelia (**Fig. 1**). However, calyces vary in their shape in different regions of amniote vestibular epithelia (Lysakowski et al. 2011), and shorter “proto-calyces” are reported to exist in fish and amphibians (Honrubia et al. 1989; Lanford and Popper 1996; Lysakowski and Goldberg 2004; Boyle et al. 2018). To address the effects of calyx geometry on the two components (*ϕ_SC_* and [*K*^+^]_*SC*_) of NQT, we studied both the steady-state condition and the step response (**Fig. 7**) for different calyx heights, while retaining the same ion channel and transporter expression per unit area of membrane. As the calyx was truncated (**Fig. 7**, ‘1/2’, ‘1/4’) relative to the standard geometry (**Figs. 1** and **7**, ‘Calyx’), both *ϕ_SC_* and [*K*^+^] _*SC*_ were closer to perilymph values (0 mV and 5 mM, respectively) at rest (**Fig. 7A,D**), had smaller responses to stimuli (**Fig. 7B,E**) and no spikes were initiated in the afferent fiber. However, the model does suggest that these truncated calyces, which resemble proto-calyces of anamniotes (assuming a similar distribution of ion channels) could accumulate K^+^, and experience noticeable changes in *E_K_* (**Fig. 7F**) that facilitate afferent depolarization. This effect on *ϕ_C_*, at the onset of bundle stimulation, is small, ~2 mV (**Fig. 7C**). On the other hand, elongation of the calyx height by 2 μm (**Fig. 7**, ‘Ext’) increased *ϕ_SC_* and [*K*^+^_*SC*_ at rest, increased rate and magnitude of stimulus-evoked changes in *ϕ_C_*, and reduced transmission latency (**Fig. 7C**). This greater height is found in the calyces of “dimorphic” afferents (Lysakowski et al. 2011), which have both calyx and bouton endings.

**Fig. 7.**
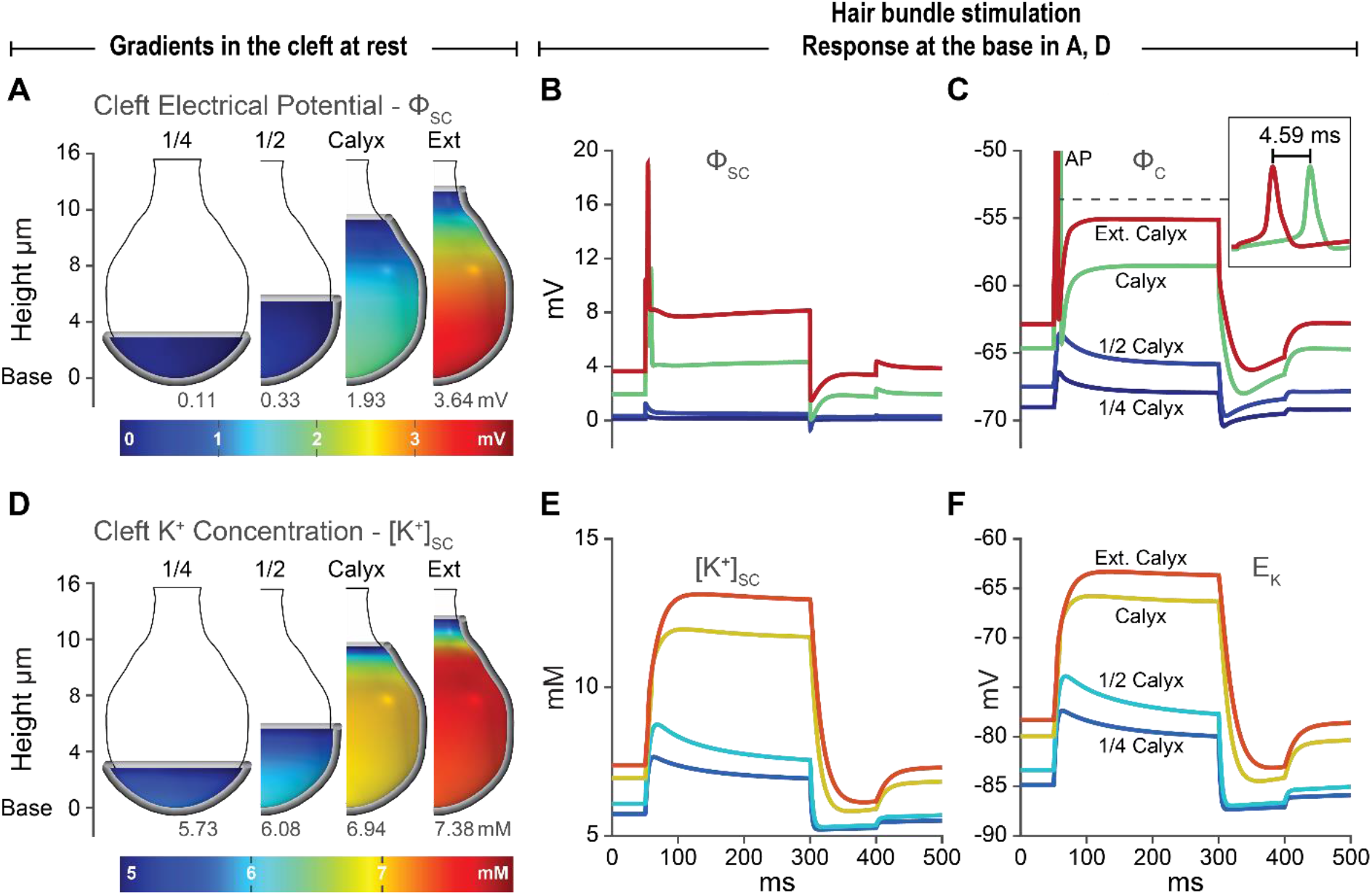
Magnitude and speed of nonquantal transmission increase with calyx height. **A, D,** The 3D visualizations represent a shell the width of the cleft space resting between the hair cell and calyx. Silver edges delineate calyces of different heights. Spatial gradients of *ϕ_SC_* and [*K*^+^]_*SC*_ are visible within these regions. The outer surface of the hair cell above the calyx is exposed to perilymph (ϕ = 0 mV, ground and [K^+^] = 5 mM). Values at the base of the synapse are shown above the color bars. **B, C,E,F,** Changes in *ϕ_SC_*, *ϕ_C_* [*K*^+^]_*SC*_ and E_K_ increase with calyx height. Curve colors correspond to those at the base of each calyx in A and D at steady state. **A,** The spatial distribution of *ϕ_SC_* at rest as a function of calyx height. **B,** *ϕ_SC_* at the base of the synapse over the course of step bundle displacement (same as in Fig. 2A). **C,** Changes in *ϕ_C_* (the transmitted signal). The inset demonstrates the reduction in time to spike as calyx height is increased by 2 *μm* from 11.3 to 13.3 *μm.* **D,** The spatial distribution of [*K*^+^]_*SC*_ at rest. **E,** [K^+^Lc at the base of the synapse over the course of step bundle displacement. **F,** *E_K_* values corresponding to changes in [*K*^+^]_*SC*_. Compare with *ϕ_SC_* (**B**).

**Fig. 7** shows that increasing calyx height (and thereby extending the synaptic cleft) increases the resting values of *ϕ_SC_* and [*K*^+^]_*SC*_, depolarizes both hair cell and calyx potentials, opens more ion channels at rest, increases stimulus-evoked changes in both *ϕ_SC_* and [*K*^+^]_*SC*_, increases the likelihood of post-synaptic spiking and reduces first-spike latency (**Fig. 7C**; the occurrence of a calyx spike is evident in the *ϕ_SC_* traces for the standard and extended calyx). Simulations of calyx heights at finer intervals indicated that NQT-initiated spiking required a calyx height within 95% of the standard geometry (11.3 μm), suggesting the central calyx height may reflect a minimal value required for NQT-driven spiking. Interestingly, once this minimal length is achieved, further increases in calyx length have a significant effect on the magnitude and rate of NQT and time-to-spike in the calyx. In addition to an increase in the calyceal membrane, this reflects an increase in the extent of the synaptic cleft and the associated axial resistance and distance to perilymph.

## Discussion

We have constructed a biophysical model that incorporates ultrastructural data on the geometry of the VHCC synapse, immunocytochemical data on the expression of ion channels and transporters and electrophysiological data on kinetics of ion channels. Our goal was to understand how currents and driving forces interact to accomplish NQT at the unique VHCC synapse, which is characterized by its extensive and unfenestrated synaptic cleft, and the many low-voltage-activated potassium (KLV) channels which populate the pre- and post-synaptic membranes. The model enables the prediction of experimentally inaccessible parameters, including potassium concentration [*K*^+^]_*SC*_ and electrical potential *ϕ_SC_* in the synaptic cleft. The model also enables us to track dynamic changes in these parameters in response to hair bundle deflection and paired voltage clamp conditions. These capabilities have produced results which explain experimental recordings of both fast (Songer and Eatock 2013; Contini et al. 2020) and slow (Highstein et al. 2014; Contini et al. 2017) NQT. In the process, we were able to identify three overlooked features in existing experimental recordings which support the existence of an electrical potential in the synaptic cleft and its role in NQT as predicted by the model: the frequency dependence of the phase of NQT (Songer and Eatock 2013); the fast retrograde events in hair cell potential during retrograde transmission of post-synaptic spikes (Contini et al. 2017); and the shape of fast post-synaptic currents (Contini et al. 2020).

The VHCC model permits a clear description of the sequence of biophysical events that give rise to NQT, summarized in **Fig. 8** and shown in **Figs. 2,3**. Modulation of hair cell potential immediately changes the synaptic cleft potential *ϕ_SC_* and more gradually changes the potassium potential *E_K_.* Both are accomplished by change in a large K^+^ current through the non-inactivating, low-voltage-activated potassium conductance, g_K,L_, on the hair cell basolateral membrane. The properties of g_K,L_ suit its role in NQT. First, g_K,L_ is >50% open at the hair cell resting potential, allowing the receptor potential to immediately alter membrane current and thereby *ϕ_SC_*. Second, g_K,L_ is non-inactivating, ensuring that significant K^+^ will accumulate in the cleft during sustained or low-frequency hair cell stimulation. The changes in cleft electrical and K^+^ potentials alter the driving forces of currents across the calyx inner face. The calyx inner face contains a large number of different low-voltage-activated channels (Kv7), which are also non-inactivating and significantly open at rest (~30%). Upon stimulation, increases in *ϕ_SC_* and [*K*^+^]_*SC*_ reduce the outward Kv7 current and depolarize the calyx. In short, the presence of open K_LV_ channels in the pre- and post-synaptic membranes enables NQT as previously proposed (Eatock 2018; Spaiardi et al. 2020). The VHCC model clarifies the role of *ϕ_SC_* and *E_K_* as driving forces for currents through all channels facing the synaptic cleft.

**Fig. 8.**
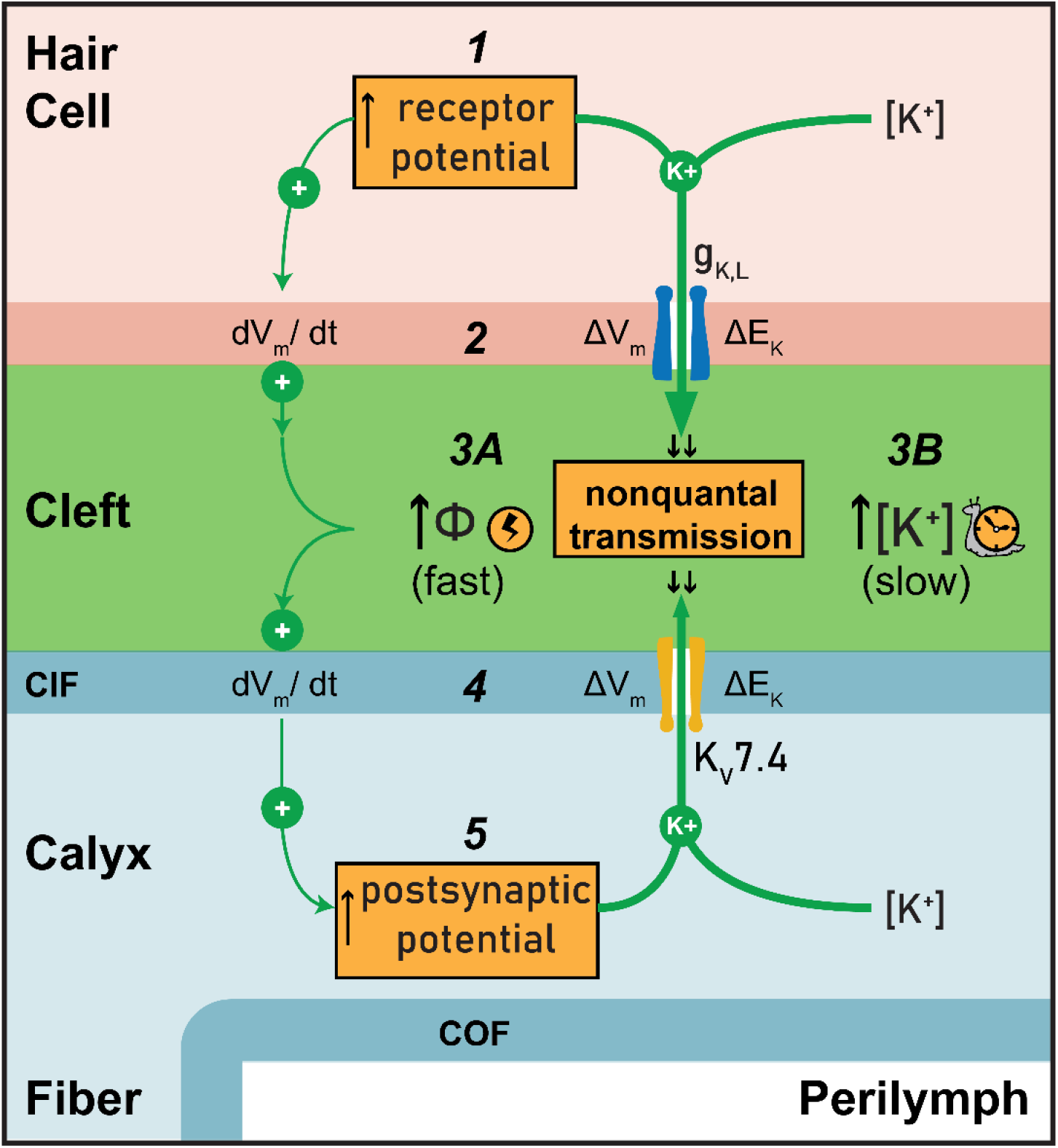
Nonquantal Transmission. 1. Depolarizing receptor potential due to positive bundle deflection.
2. Change in hair cell membrane voltage drives capacitive currents (left pathway) and *increases* outward resistive current through g_K,L_ (right pathway).
3. Changes in the synaptic cleft
  a. Fast rise in potential in the synaptic cleft, *ϕ_SC_*
  b. Gradual rise in cleft potassium concentration [*K*^+^]_*SC*_
4. Changes in *ϕ_SC_* and [*K*^+^]_*SC*_ alter membrane voltage and E_K_ respectively across the calyx inner face. This drives capacitive currents (left pathway) and *decreases* outward resistive current through K_V_7.4 (right pathway). During bundle stimulation, resistive currents in steps 2,4 are much larger than capacitive currents.
5. The postsynaptic potential depolarizes (Calyx is depolarized). Note that the cleft compartmentalizes the calyx membrane into inner (CIF) and outer (COF) faces, with different trans-membrane voltages. The postsynaptic potential corresponds to that measured with respect to perilymph and across the calyx outer face.

The hypothesis that currents flow through open channels on either side of the membrane was tested experimentally and qualitatively presented as ‘resistive coupling’ between presynaptic g_K,L_ and postsynaptic HCN channels (Contini et al. 2020). Fast post-synaptic currents were presented as evidence for resistive coupling but the driving forces that shaped these currents were not understood, and some were thought to be artefacts. Using the VHCC model, we have demonstrated that changes in cleft electrical potential, shaped by capacitive and resistive currents from the hair cell, underlie these fast post-synaptic currents.

At the calyx of Held, a different large synapse important for auditory processing (Borst and Soria van Hoeve 2012), analysis of ephaptic coupling (Sierksma and Borst 2021) suggested that although pre-synaptic capacitive and resistive currents occur, the fenestrated morphology of the calyx of Held serves to mitigate synaptic cleft potentials to ensure presynaptic calcium channels sense the full presynaptic action potential and rapidly actuate neurotransmitter release. In contrast, what naturally emerges from the VHCC model is that the unfenestrated morphology of the VHCC synapse is specialized to actively use cleft [*K*^+^]_*SC*_ and *ϕ_SC_* as complementary mechanisms that mediate NQT and benefit signal transmission between the sensory hair cell and the afferent neuron. The slower changes in [*K*^+^]_*SC*_ increase the gain of NQT for slow or sustained hair bundle deflections, but do not alone explain the observed phase response. The near-instantaneous change in *ϕ_SC_* enables the calyx to closely follow changes in hair cell potential and explains the short latency and phase characteristics of NQT observed experimentally (Songer and Eatock 2013; Contini et al. 2020). Ultimately NQT also enables afferent neurons to follow the stimulus up to relatively high frequencies. The calyx geometry is essential for both fast and slow NQT; as calyx height increases the efficacy of both mechanisms, *ϕ_SC_* and [*K*^+^]_*SC*_, increases and transmission latency decreases.

These findings explain the exceptional speed of neurotransmission between type I hair cells and vestibular afferents, which is even faster than the quantal transmission between cochlear hair cells and afferents (McCue and Guinan 1994). At its fastest, quantal transmission imposes a 0.4-0.6-ms delay (Lisman et al. 2007), reflecting the time required for multiple steps from the opening of presynaptic voltage-gated calcium channels to the opening of postsynaptic neurotransmitter-gated receptor channels. For fast vestibular reflexes, with latencies as short as 5 ms between head motion and motor response (Huterer and Cullen 2002), overcoming quantal transmission delay and providing an additional mode of communication through NQT at the first synapse in the pathway is a significant advantage. Our results indicate that robust NQT, enabled by the evolution of the full calyx, substantially reduces the latency and advances the phase of vestibular signals that drive neural circuits controlling gaze, balance, and orientation.

In conclusion, the VHCC model provides a quantitative description of NQT and describes two parallel mechanisms of NQT: changes in the cleft electrical potential and changes in the potassium potential. It also delineates the roles of specific ion channels and calyx geometry in NQT. Here we have focused on a simple calyx from central or striolar zones of vestibular epithelia; the model can be extended to explore the functional effects of other afferent terminal forms, such as the complex calyces that enwrap multiple type I hair cells and the elaborate dimorphic terminal arbors that include calyces and boutons (reviewed in Goldberg 2000; Eatock and Songer 2011).

## Data Availability

The model was implemented in Comsol Multiphysics 5.6 software. Equations and data sources are available in **SI Computational Methods and Experimental Methods.** The Comsol file (.mph) containing the model will be publicly available on Github (https://github.com/acgsci/vestibularHairCellCalyxNQT).

## Author Contributions

RAE and RMR conceived and supervised the work. ACG expanded an initial model developed by IQ. ACG and IQ wrote COMSOL and Matlab code for model implementation. RAE and AL supervised the acquisition of data presented in the Supplement. The manuscript was written by ACG, RMR and RAE and revised and edited by all authors.

## Acknowledgements

This study was supported by National Institutes of Health (R01 DC012347 and DC002290) and a seed grant from the RICE ENRICH program. We are grateful to Hannah Martin for stimulating discussions and Selina Baeza-Loya for providing activation and inactivation time constants of afferent voltage-gated sodium channels.

## Supplementary Information

### Computational Methods

#### Overview

The Vestibular Hair Cell-Calyx (VHCC) model incorporates information on synapse morphology, ion transporters and ion channels expressed in type I hair cells (channel types: transduction (MET), K_V_, Ca_V_, HCN; transporter: Na^+^/K^+^-ATPase), and in calyces and afferent fibers (channel types: K_V_, Na_V_, HCN; transporters: KCC, Na^+^/K^+^-ATPase) (see **Fig. 1** in main text). The dynamic behavior of the system is determined from measured or estimated channel open probabilities, time constants and whole-cell conductances, and from assumptions (which we describe) about the activity of ion transporters based on literature. We implemented the VHCC model with finite-element-analysis simulation software, using equations for K^+^ and Na^+^ electro-diffusion in the cleft, Hodgkin-Huxley-style equations to represent voltage-dependent ion flow through ion channels, and the cable equation for electrical propagation. For each simulation, steady-state conditions are established and followed by a step or sinusoidal deflection of the hair bundle or voltage step; outputs include the basal-to-apical gradients within the synaptic cleft for K^+^, Na^+^ and electrical potential, and voltage across the hair-cell-facing postsynaptic membrane of the calyx (“calyx inner face”), and in the hemi-node and first two full nodes of the distal branch of the bipolar vestibular afferent neuron. The distal branch connects the synaptic terminals on hair cells (in this case a single calyx) to the neuronal cell body in the vestibular ganglion; the remainder of the distal branch, the cell body and the central branch projecting to the brain are not included in the model. Negative currents represent the flow of cations into the compartment bearing the channel or transporter in its membrane. The hair cell is assumed to be equipotential based on its small overall size, compact form and low intracellular resistance. Details of model development are described below.

#### Model Geometry

The model was designed with the COMSOL Multiphysics 5.6 software package. We used a 2D axisymmetric finite element boundary to represent the hair cell, synaptic cleft, and afferent calyx and a 1D finite element line to represent the afferent fiber (**Fig. S1**). The 2D boundary used to represent the hair cell (H), synaptic cleft (SC), the calyx inner face (CIF) and calyx outer face (COF) is an interpolation curve generated from (R,Z) coordinates of the perimeter of a central-zone hair cell and calyx from the central zone of the sensory epithelium of a crista in an adult Long-Evans rat (shown in Fig. 2C of Lysakowski et al. 2011; similar calyces from the striolar zone of the rat utricle are shown in our **Fig. 1A**). The dimensions and interpolated points used to create the representative geometry in Comsol 5.6 are shown in **Fig. S1.** HN, N and M indicate the hemi-node, nodes and myelinated sections (inter-nodes) of the afferent fiber, respectively. In both 2D (hair cell and calyx) and 1D (fiber) geometries, local properties such as individual channel conductance density, time constants, and steady state expressions for channel gating variables were defined. Note that the apex of the calyx occurred at 11.3 μm relative to the calyx base for all simulations in the manuscript, except for the variations in calyx height presented in **Fig. 7**.

**Fig. S1.**
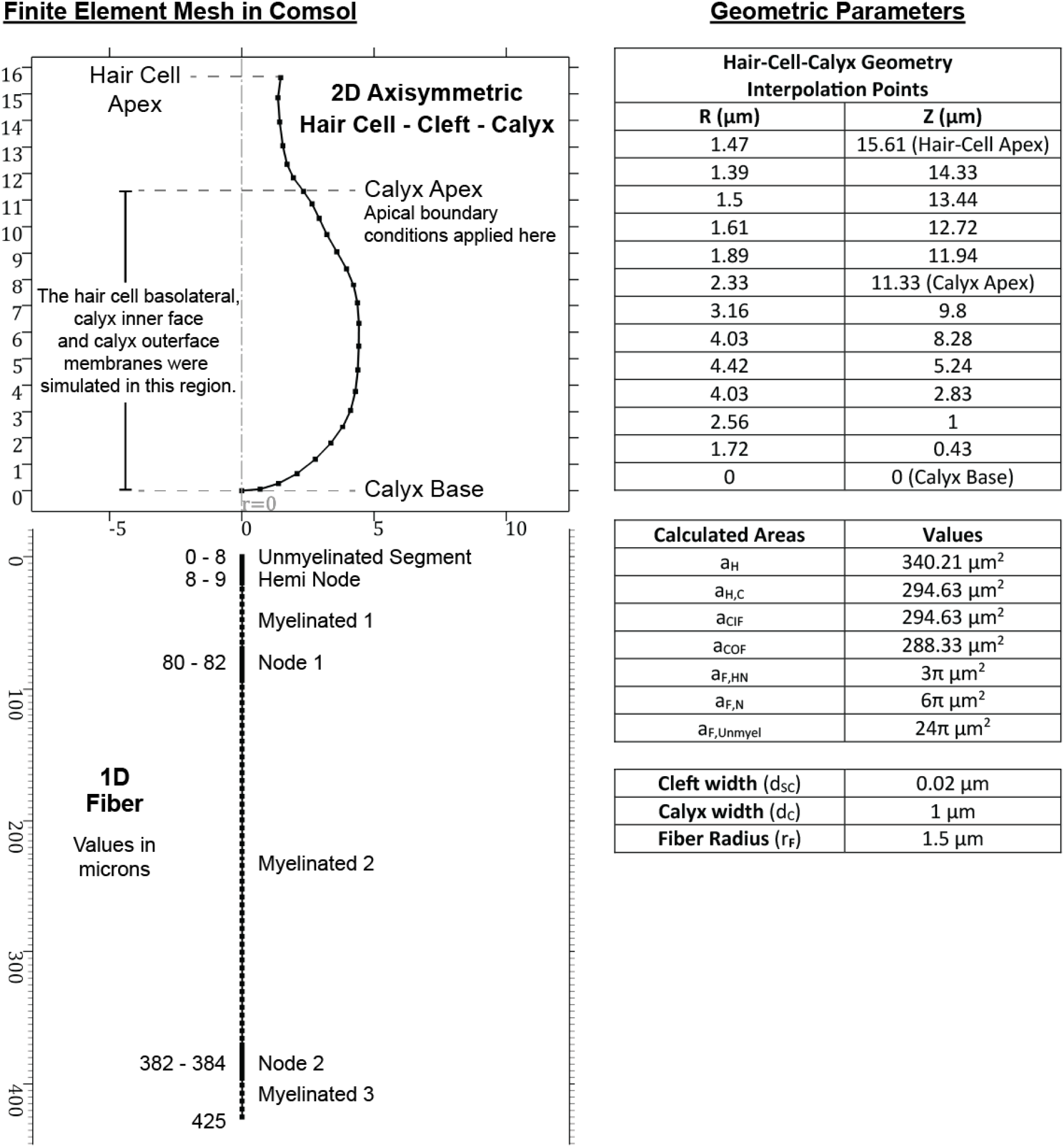
Hair-Cell-Calyx and Fiber Geometries in the COMSOL 5.6 GUI. The 2D axisymmetric mesh representing the hair cell, synaptic cleft and calyx (top) was modelled based on the geometry of a type I hair-cell-calyx synapse from central/striolar zones of rodent vestibular organs. Dots in the above geometry represent finite element nodes, and lines represent connecting edge elements. The interpolation points were based upon the geometry of a hair cell and calyx reported in Fig. 2C of Lysakowski et al. (2011). *d_C_* was obtained from the same calyx. *d_SC_* was obtained from Spoendlin (1966). The fiber radius (*r_f_*) is based on experimental measurements (Lysakowski et al. 1995; Hoffman and Honrubia 2002). The fiber end was treated as a closed boundary. Location of the hemi-node and the first node were estimated from confocal micrographs (Lysakowski et al. 2011). *α_H,C_* denotes the surface area of the hair cell within the calyx.

**Fig. S2.**
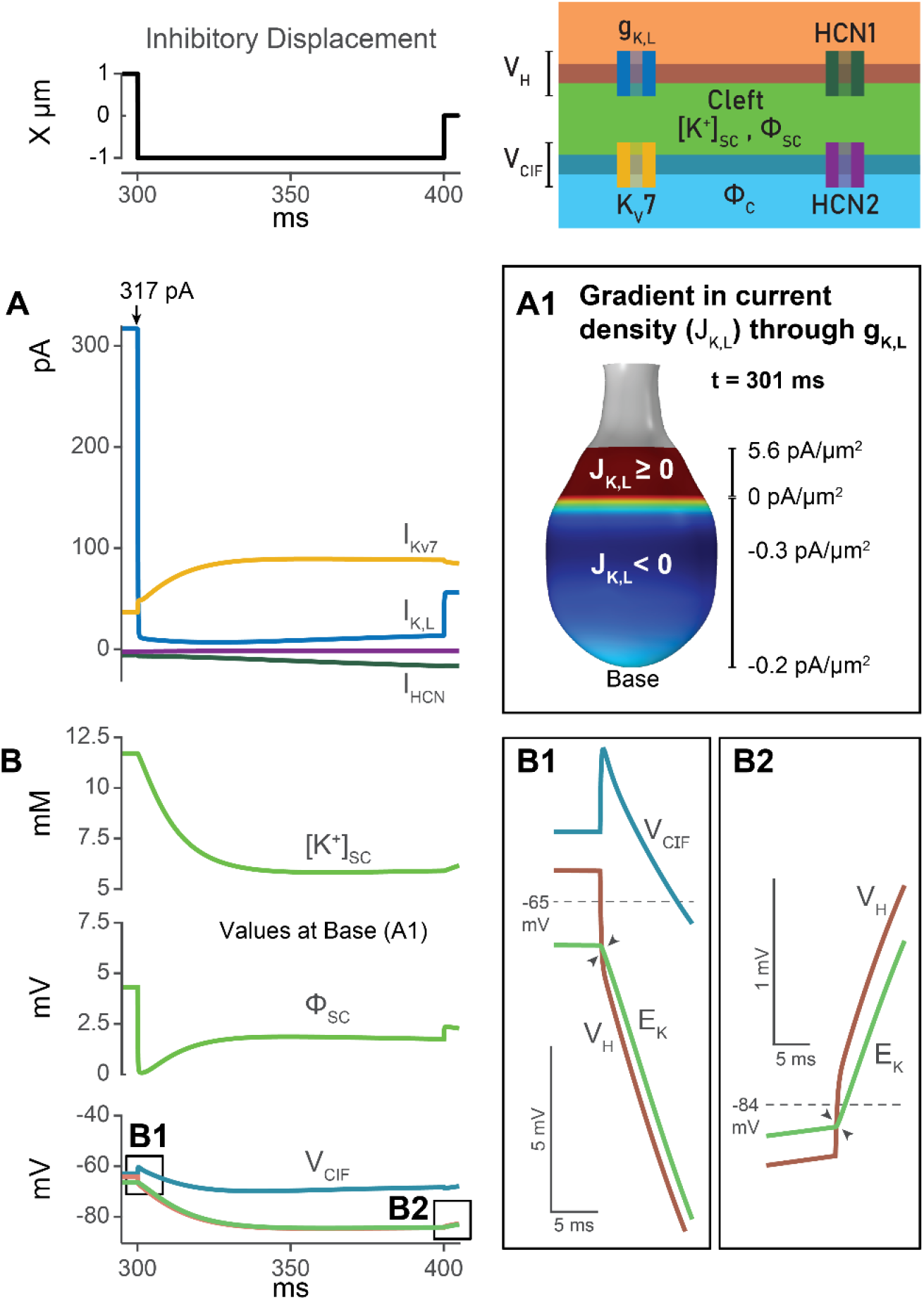
Gradients in current density exist from base to apex of the synaptic cleft. **A,** Net currents through g_K,L_ (hair cell, blue), K_V_7 (calyx, gold) and HCN channels (hair cell, dark green, and calyx, purple) following the onset of the full negative step. **A1,** Gradient in current density through g_K,L_ 1 ms after the negative step at 300 ms. Small currents flow into the hair cell in basal regions, and larger currents flow out of the hair cell near the apical regions of the synaptic cleft. The direction and magnitude of the current results from the difference between V_H_ and E_K_ which varies from base to apex and with the ongoing stimulus. This behavior is illustrated at the base of the synapse in B, B1 and B2. **B,** [K^+^]_SC_, ϕ_SC_, V_H_, V_CIF_ and E_K_ at the base of the synapse during inhibitory stimuli. Note the crossover of V_H_ and E_K_ following the onset (**B1**) and the termination (**B2**) of the negative step. As the driving force for currents is (V_H_ - E_K_), the separation between the V_H_ and E_K_ curves determines current magnitude and current direction is outward if V_H_ is positive to E_K_. During the negative step current through g_K,L_ in the basal regions in A1 are inward because V_H_ is negative to E_K_.

**Table S1.** Abbreviations.

|  |  |
| --- | --- |
| **$H, C, SC, CIF, COF, F$** | Subscripts denote the hair cell ($H$), calyx ($C$), synaptic cleft ($SC$), calyx inner face ($CIF$), calyx outer face ($COF$) and fiber ($F$) respectively. |
| **$\phi_H, \phi_C, \phi_{SC}, \phi_F$** | Electrical potential in the hair cell, calyx, synaptic cleft and fiber respectively. |
| **$\rho_F$** | Resistivity of the afferent fiber. $1/\sigma_F$ |
| **$[K^+]_{SC}, [Na^+]_{SC}$** | $K^+$, $Na^+$ concentrations in the synaptic cleft |
| **$c_H$** | Total hair cell capacitance |
| **$C_m$** | Specific membrane capacitance |
| **$Ca_v, K_v, Na_v$** | Voltage-gated calcium, potassium and sodium channels |
| **$D_K, D_{Na}$** | Effective diffusion coefficients for $K^+$, $Na^+$ in the synaptic cleft |
| **$d_{SC}$** | Width of the synaptic cleft |
| **$g_{x,max}$** | Maximum whole-cell conductance of specific channel type |
| **$G_{x,max}$** | Maximum conductance density of specific channel type |
| **$g_{K,L}$** | Low-voltage-activated potassium conductance of type I hair cells |
| **$HCN$** | Hyperpolarization-activated nucleotide-gated (channels) |
| **$I_{MET}$** | Mechanotransduction current |
| **$J_x$** | Current density through channel or transporter 'x' |
| **$J_{M_x}$** | Current density through membrane 'x' |
| **$KCC$** | Potassium Chloride Cotransporter |
| **$K_v7$** | Voltage-gated potassium channels formed by subunits encoded by the KCNQ gene family |
| **$P_{open}$** | Channel open probability |
| **$VHCC$** | Vestibular Hair Cell-Calyx |
| **$V_m$** | Membrane voltage |

**Table S2.** Equilibrium Potentials and Voltages Across Membranes.

| Property |  |  | Hair Cell (H) | Calyx inner face (CIF) | Calyx outer face (COF) | Fiber (F)<br>HN\|N\|M |
| --- | --- | --- | --- | --- | --- | --- |
| $E_K$ | $K^+$<br>Equilibrium Potential | $mV$ | $26 \cdot \ln \left( \frac{[K^+]_{sc}}{150} \right)$ | $26 \cdot \ln \left( \frac{[K^+]_{sc}}{150} \right)$ | $26 \cdot \ln \left( \frac{[K^+]_{sc}}{150} \right) = -88.4$ | $-88.4$ |
| $E_{Na}$ | $Na^+$<br>Equilibrium Potential | $mV$ | $26 \cdot \ln \left( \frac{[Na^+]_{sc}}{12} \right)$ | $26 \cdot \ln \left( \frac{[Na^+]_{sc}}{12} \right)$ | $26 \cdot \ln \left( \frac{140}{12} \right) = 63.8$ | $63.8$ |
| $V_m$ | Membrane Voltage | $mV$ | $\phi_H - \phi_{sc}$ | $\phi_C - \phi_{sc}$ | $\phi_C$ | $\phi_F$ |

### General Modeling of Membrane Currents

Membrane channels and transporters were distributed in the various compartments based on experimental observations (**Table S6, S7**). Abbreviations used for ion channels and other parameters are listed in **Table S1**. For each type of ion, local current density through a particular channel type ‘x’ was described by **Eq. S1**:

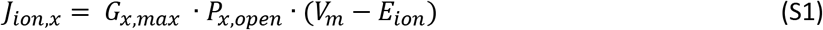

*G_x,max_* is the maximum conductance density through a channel ‘x’ per unit area of membrane, obtained by dividing the maximum whole-cell conductance, *g_x,max_*, by the surface area of the relevant membrane. *P_x,open_*, calculated as a function of channel gating variables using the Hodgkin-Huxley formalism, is the percentage of the maximal conductance available. The equations describing the kinetics of channel gating variables as functions of membrane voltage and current densities through each channel type are available in **Table S6**. *V_m_* is the transmembrane potential as defined in **Table S2** for each membrane in the model. *E_ion_* is the equilibrium potential of the ion species across the membrane and is continuously updated as the ion concentration changes in the synaptic cleft. In the notation for current densities, K, Na, O, indicate the potassium, sodium, and other ions such as Ca^2+^ or Cl^-^. Current densities and fluxes, *J*, through channels and transporters were summed to describe the current densities of ionic species across Hair cell (H) and Calyx Inner Face (CIF) or Calyx Outer Face (COF) membranes. The species currents were in turn summed to obtain the total current densities through each membrane (*J_MH_*, *J_M_CIF__*, *J_M_COF__*, *J_M_F__*) (**Table S3**).

See **Table S6** for equations of current densities and kinetics for each channel type.

**Table S3.** Local and Net Current Densities Across Individual Membranes.

|  |  |  |  |  |  |
| --- | --- | --- | --- | --- | --- |
| $J_{KH}$ | $J_{K,L} + J_{K,HCN1} + J_{K,NaK}$ | $J_{NaCIF}$ | $J_{Na,HCN2} + J_{Na,NaK}$ | $J_{MH}$ | $J_{KH} + J_{NaH} + J_{OH}$ |
| $J_{NaH}$ | $J_{Na,HCN1} + J_{Na,NaK}$ | $J_{OCIF}$ | $-J_{KCC4}$ | $J_{MCIF}$ | $J_{KCIF} + J_{NaCIF} + J_{OCIF}$ |
| $J_{OH}$ | $J_{Ca}$ | $J_{KCOF}$ | $J_{K,HCN2} + J_{K,NaK}$ | $J_{MCOF}$ | $J_{KCOF} + J_{NaCOF} + J_{Leak}$ |
| $J_{KCIF}$ | $J_{KV7.4} + J_{KV11.1} + J_{K,HCN2} + J_{K,NaK} + J_{KCC4}$ | $J_{NaCOF}$ | $J_{Na,HCN2} + J_{Na,NaK}$ | $J_{MF}$ | $J_{KV7.x} + J_{KV1.x} + J_{KV3.4} + J_{NaV1.x}$ |

### Governing Equations

Continuity equations describe the electric potential and ion concentrations in the various compartments.

#### Hair Cell Electrical Potential (ϕ_H_)

The time dependence of the hair cell electrical potential is calculated as a function of hair cell capacitance (*C_H_*), mechanotransduction current (*I_MET_*), and current across the basolateral membrane:

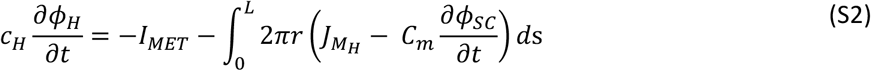

*I_MET_* is calculated according to an instantaneous current-displacement relationship (Holt et al. 1997) and a maximal conductance of 5 nS, based on ~50 transduction channels of ~100-pS single-channel conductance (Corey et al. 2019). We calculated the total capacitance of the hair cell (*C_H_* = 6.4 *pF*) by multiplying the surface area of the cell by the specific capacitance of the membrane (*C_m_*).

#### Concentration and Electro-diffusion in the Synaptic Cleft

[*K*^+^]_*SC*_ and [*Na*^+^]_*SC*_ were calculated with the Nernst-Planck electro-diffusion equation for each ion:

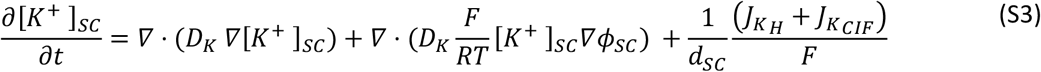

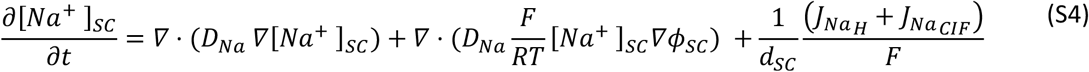

*D_K_,* the effective diffusion coefficient for *K*^+^, is calculated by dividing the diffusion coefficient of *K*^+^ in aqueous solution at 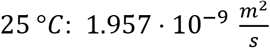 (Lide 2005) by the square of the tortuosity coefficient 1.55 (Nicholson and Phillips 1981), 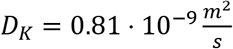. Similarly, 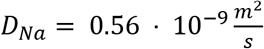. The volume of the synaptic cleft is assumed to be constant and the width (*d_SC_*) and length of the synaptic cleft are fixed.

#### Electrical Potential in the Synaptic Cleft (*ϕ_SC_*)

At bouton synapses, the electrical potential in the synaptic cleft, *ϕ_SC_*, is assumed to be that of the extracellular medium (at reference potential or ground, ~0 mV) because the resistance of the cleft is assumed to be low. At the VHCC synapse, in contrast, the extended synaptic cleft created by the calyx geometry poses an increased resistance and large K^+^ currents that flow during transduction may allow a significant cleft potential. *ϕ_SC_* was calculated following the law of charge conservation as a function of the resistive and capacitive currents across both pre-synaptic (hair cell) and post-synaptic (calyx) membranes, as well as electro-diffusion of ions:

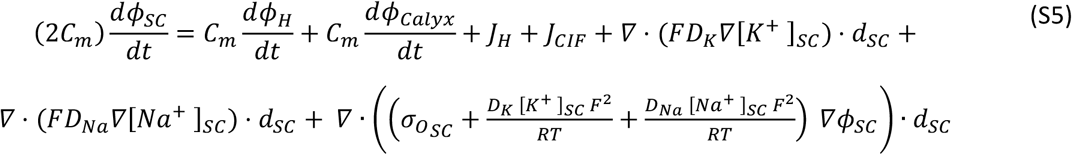

At room temperature, the terms 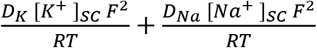resolve to ~ 300 - 400 nS/μm when [*K*^+^]_*SC*_ and [*Na*^+^]_*SC*_ range between 5-40 and 120-140 mM, respectively. The total conductivity contribution from ions (*σ_O_SC__*) other than K^+^ and Na^+^ is set equal to 600 nS/μm to be consistent with the 1000 nS/μm used in the calyx and afferent fiber. 1000 nS/μm (equivalent to 1 S/m) is within the conductivity range (0.75 – 1.45 S/m) reported from measurements in tumor cells (Wang et al. 2017), hippocampal neurons (Zhou et al. 2016) and cerebrospinal fluid (Baumann et al. 1997). The diffusion of other ions, e.g., chloride, was not explicitly considered and their concentrations within the synaptic cleft are assumed to be constant and a part of the *σ_O_SC__* term.

#### Electrical Potential in the Calyx (*ϕ_C_*)

The electrical potential within the calyx was calculated as a function of the resistive and capacitive currents across both the post-synaptic membrane and the calyx outer face membranes as well as the longitudinal current from the afferent fiber.

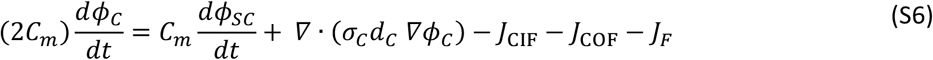

The current from the fiber to the calyx, *J_F_*, is equal to 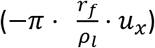, where *r_f_* the radius of the fiber is 1.5 μm, *ρ_l_* the longitudinal resistivity is 1 MΩ μm, and *u_x_* is the longitudinal potential gradient within the fiber. The fiber radius used is comparable to diameters reported for calyx afferents (Hoffman and Honrubia 2002).

#### Electrical Potential in the Afferent Fiber (*ϕ_F_*)

To describe the potential in the afferent fiber, *ϕ_F_,* we implemented the cable equation along the 1D line representing the fiber:

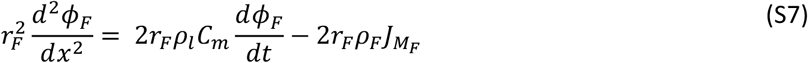

Channel gating variables and conductance densities (see **Table S6**) were defined for the hemi-node, nodes and myelinated regions of the fiber as shown in **Fig. 1**. Gating variables for channels present on the fiber were treated similarly to those on the calyx and hair cell.

#### Initial and Boundary Conditions

Before running the simulation, the initial value of *ϕ_H_* was set at −70 mV, within the normal range of reported resting potentials for rodent type I hair cells (Rennie and Correia 1994; Rennie et al. 1996; Rüsch and Eatock 1996; Chen and Eatock 2000; Lim et al. 2011; Spaiardi et al. 2017). Where necessary, initial values of *ϕ_H_* were varied ±10 mV to assist solver convergence. An initial value of 0 mV was used for the potential in the synaptic cleft, *ϕ_SC_*. The boundary condition *ϕ_SC_ =* 0 mV was applied at the apical end of the synaptic cleft. An initial value of −70 mV was used for *ϕ_Calyx_* and *ϕ_F_*, consistent with recorded resting potentials in rodent calyces ranging −60 to −70 mV in (Meredith and Rennie 2015; Sadeghi et al. 2014; Songer and Eatock 2013). The boundary condition *ϕ_Calyx_* = *ϕ_F_* was applied at the base of the calyx and the far end of the fiber was treated as a closed boundary. [*K*^+^]_*SC*_ and [*Na*^+^]_*SC*_ were given initial values of 5 mM and 140 mM, respectively, throughout the cleft, based on the external (bath) recording solutions of electrophysiological data referenced in the model and similar to perilymph facing the outer face of the calyx and other extracellular salines. K^+^ and Na^+^ concentrations were pinned at these values at the apical end of the cleft but were allowed to vary elsewhere within the cleft. To better understand Non-quantal Transmission (NQT), we altered the initial conditions to independently analyze the contributions of changes in the synaptic cleft in (1) electrical potential and (2) potassium concentration. As described below, this theoretical exercise gave greater insight into the mechanisms of NQT.

**Table S4.**
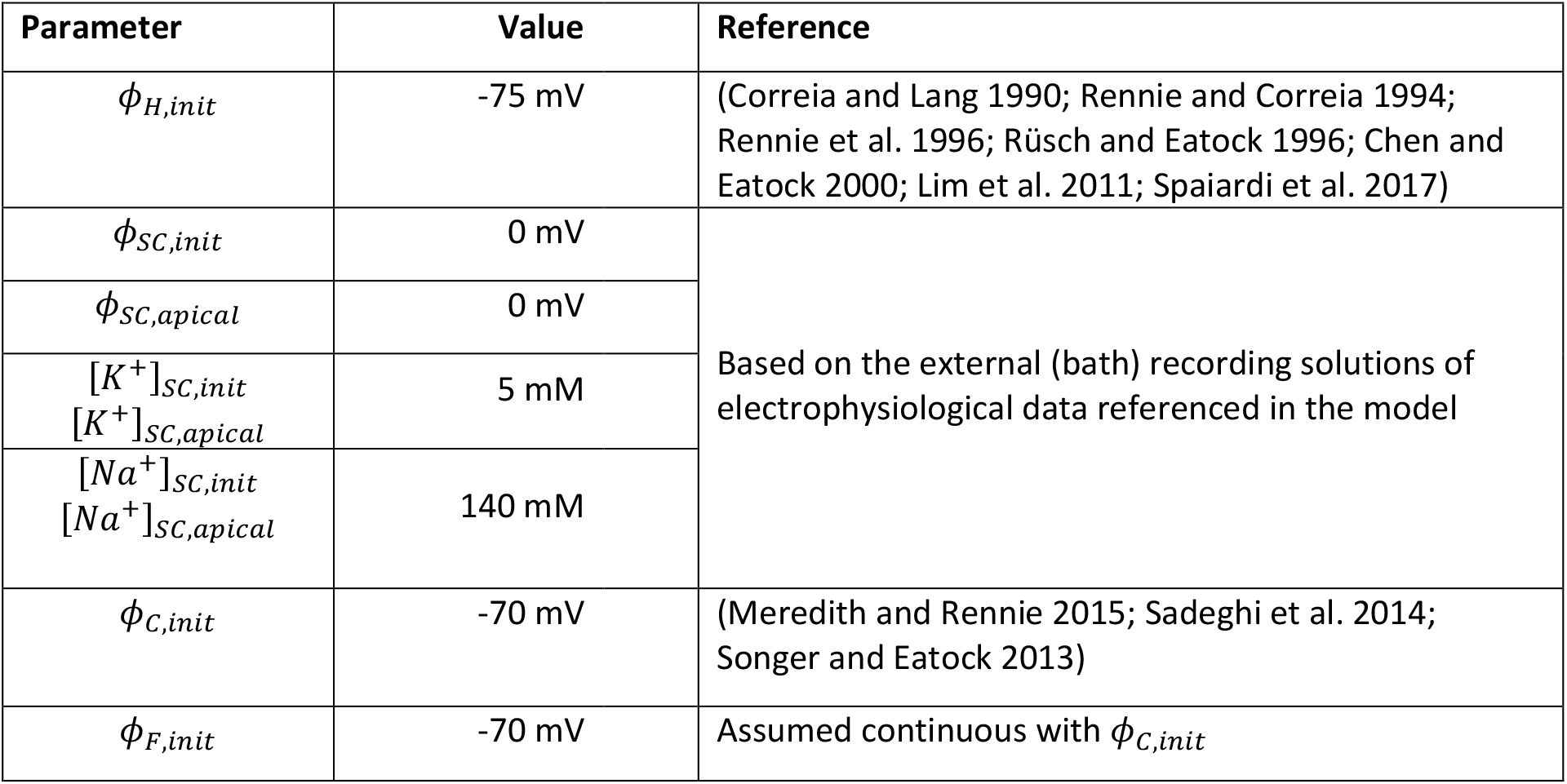
Initial and Boundary Conditions.

#### *ϕ_SC_*-Only Condition

To isolate the contribution of changes in electrical potential within the synaptic cleft to NQT, we held ion concentrations in the synaptic cleft constant at 5 mM for [*K*^+^]_*SC*_ and 140 mM [*Na*^+^]_*SC*_ (**Eqs. S3 and S4**) and removed terms for ion diffusion in the cleft from **Eq. S5**.

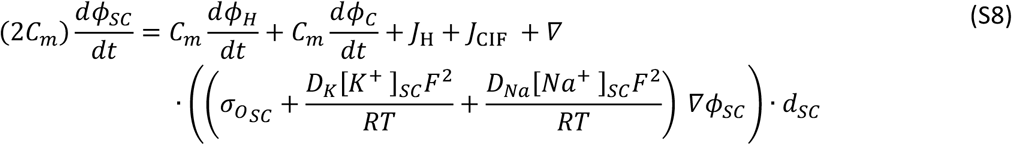

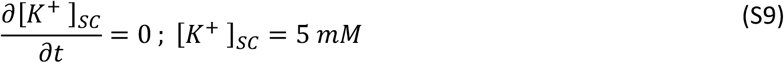

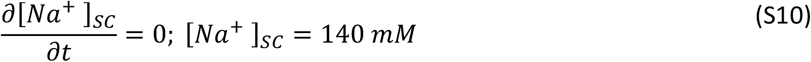

#### [*K*^+^]_*SC*_-Only Condition

To isolate the contribution of changes in K^+^ concentration in the synaptic cleft to NQT, we fixed *ϕ_SC_* at 0 mV and removed the electrical drift terms in **Eqs. S3 and S4**.

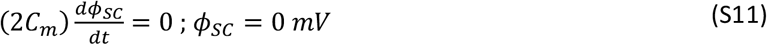

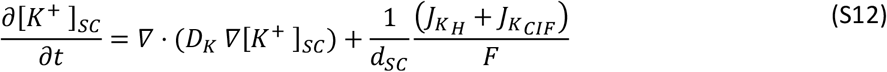

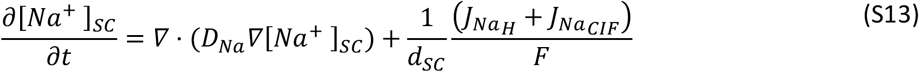

### Model Simulation and Solver Conditions

In the COMSOL 5.6 software, the finite element mesh was generated based on geometric parameters for the hair-cell-calyx and fiber (**Fig. S1**). The number and distribution of finite element nodes (hair-cell-calyx: 26 nodes, 25 edge elements; fiber: 95 nodes, 94 edge elements) was chosen to minimize error in the solutions of the governing equations. To obtain the resting conditions, we ran the model simulation with a stationary solver. For a step or sinusoidal displacement of the hair cell’s stereociliary bundle or hair cell voltage steps, we used a time-dependent solver to determine the responses in the hair cell, synaptic cleft, calyx and afferent fiber. The time-dependent solver was run using the backward difference formula (BDF) method. The temporal resolution (maximum time step) was: 1 ms for resting conditions before stimulus onset; 0.01 ms from 1 ms before stimulus onset to 10 ms after stimulus onset; and 0.1 ms for all other intervals. The model simultaneously solves **Eqs. S1-S7** to simulate the electrical potential of the hair cell and the spatiotemporal evolution of electrical potential, potassium concentration, sodium concentration in the synaptic cleft, and potential in the afferent fiber.

**Table S5.** Conductances, Diffusion Coefficients and Capacitances.

|  |  |  |
| --- | --- | --- |
| $\sigma_C$ | 1000 nS/$\mu\text{m}$ | (Baumann et al. 1997; Geddes and Baker 1967; Zhou et al. 2016; Wang et al. 2017) |
| $C_m$ | 0.01 pF/$\mu\text{m}^2$ | (Curtis and Cole 1938; Gentet et al. 2000) |
| $C_{m,Myel}$ | 0.00002 pF/$\mu\text{m}^2$ | (Stämpfli 1954) |
| $D_K$ | 0.81 $\mu\text{m}/\text{ms}$ | (Lide 2005). See methods near **Eq. S4** |
| $D_{Na}$ | 0.56 $\mu\text{m}/\text{ms}$ | (Lide 2005). See methods near **Eq. S4** |
| $g_{bundle,max}$ | 5 nS | (Holt et al. 1997) |
| $g_{Ca,max}$ | 0.5 nS | (Bao et al. 2003) |
| $g_{HCN1,max}$ | 4.2 nS | (Horwitz et al. 2011) |
| $g_{HCN2,max}$ | 3.6 nS | (Horwitz et al. 2014) |
| $g_{K,L,max}$ | 80 nS | (Spaiardi et al. 2017) |
| $g_{K_V7.4,max}$ | 24 nS | (Hurley et al. 2006) |
| $g_{K_V1.x,max}$ | 6.3 nS | (Kalluri et al. 2010) |
| $G_{K_V3.4,max}$ | 0.2 nS/$\mu\text{m}^2$ | (Chabbert et al. 2001) |
| $g_{K_V7.x,max}$ | 4.5 nS | (Kalluri et al. 2010) |
| $g_{Na_V,unitary}$ | 10 pS | (Bezánilla 1987; Kay et al. 1998) |
| $\rho_{Na_V}$ | 12,000/$\mu\text{m}^2$ | (Ritchie and Rogart 1977) |

### Channel Distribution and Kinetics

The governing equations depend upon the net current densities across the pre- and post-synaptic membranes in the model. The constitutive currents and their kinetics as used in the model are detailed in **Table S6**. All channel properties used were from electrophysiological measurements near room temperature (23 – 27°C). For a given channel, *act*_∞_(*V*) and *τ_act_*(*V*) are expressions for the dependence of the activation ‘act’ (or inactivation ‘inact’) gating variable on the membrane voltage and the corresponding time constant. *act*(*t*) is the instantaneous value of the gating variable. The expressions for *act*(*V*) and *τ_act_*(*V*) were stored in variables assigned to the VHCC and fiber geometry to make them locally available during model evaluation. The maximum whole-cell conductance value *g_x,max_* for any type of channel ‘x’ was divided by the surface area to obtain the maximum conductance density *G_x,max_* to allow us to evaluate local current densities at specific sites in the model. The transporters Na^+^/K^+^-ATPase and KCC were assumed to operate at pseudo-steady state and depend only on [*K*^+^] in the cleft. On the hair cell, basolateral channels were only distributed to the part of the membrane enclosed by the calyx.

**Table S6.**
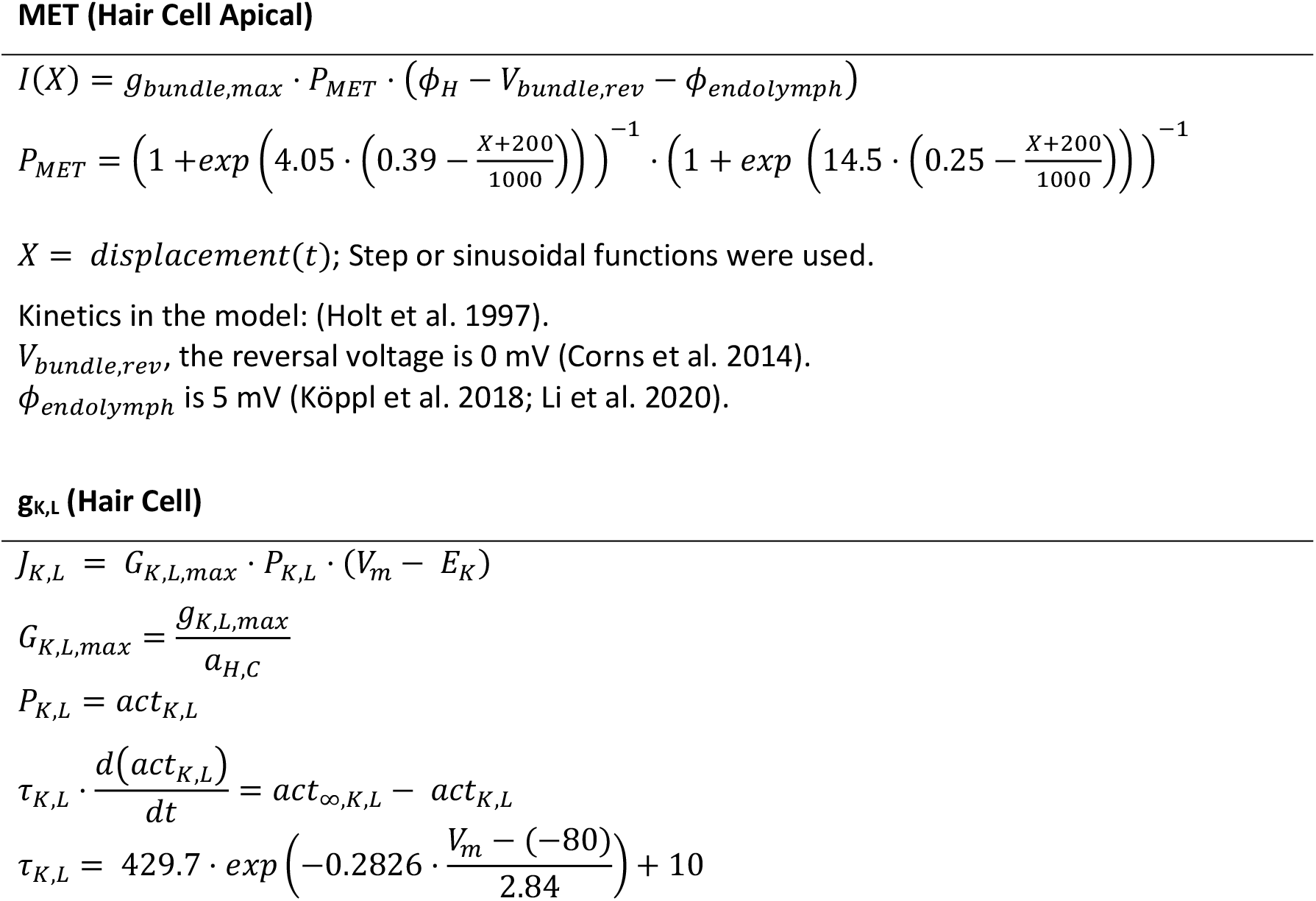
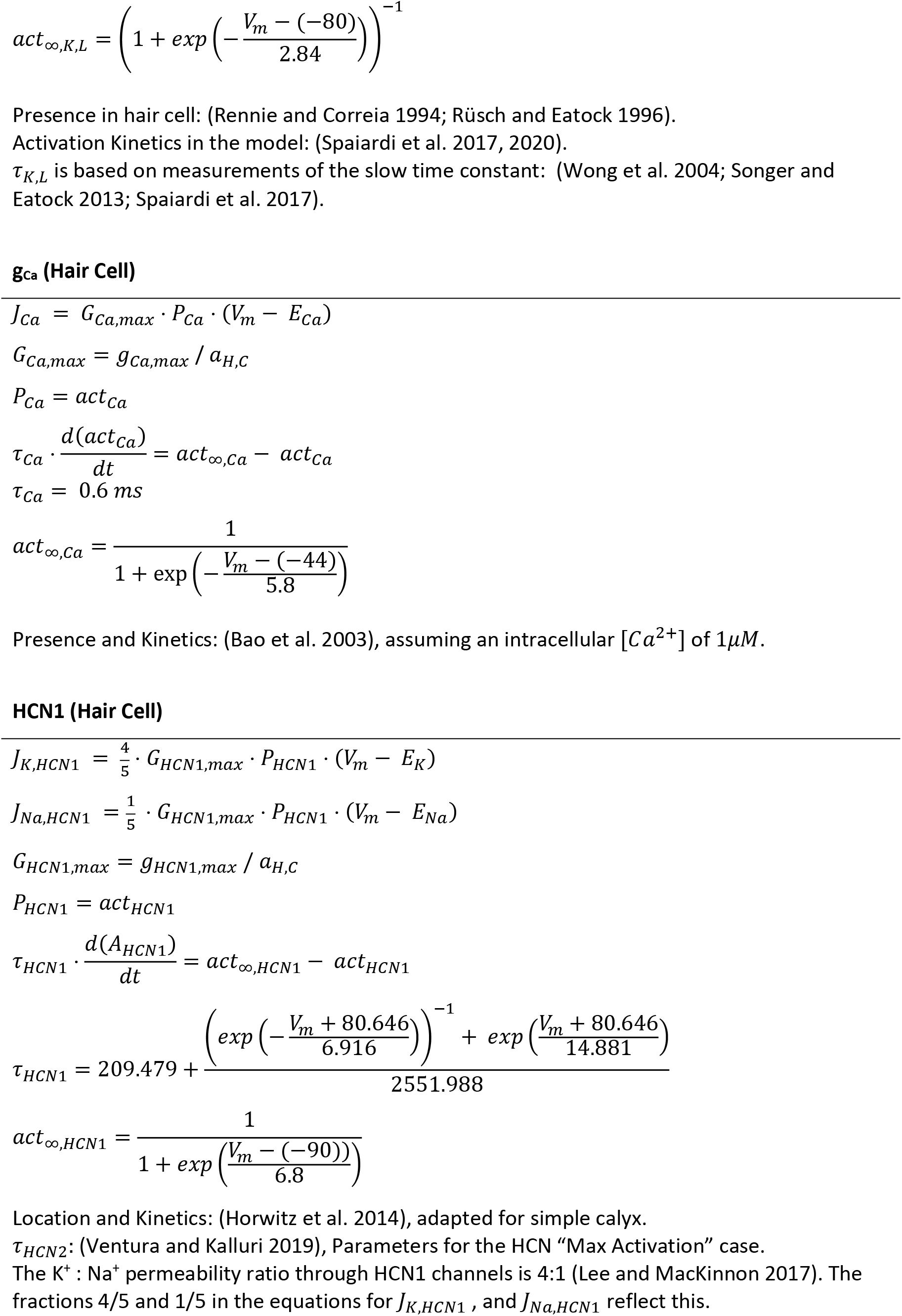
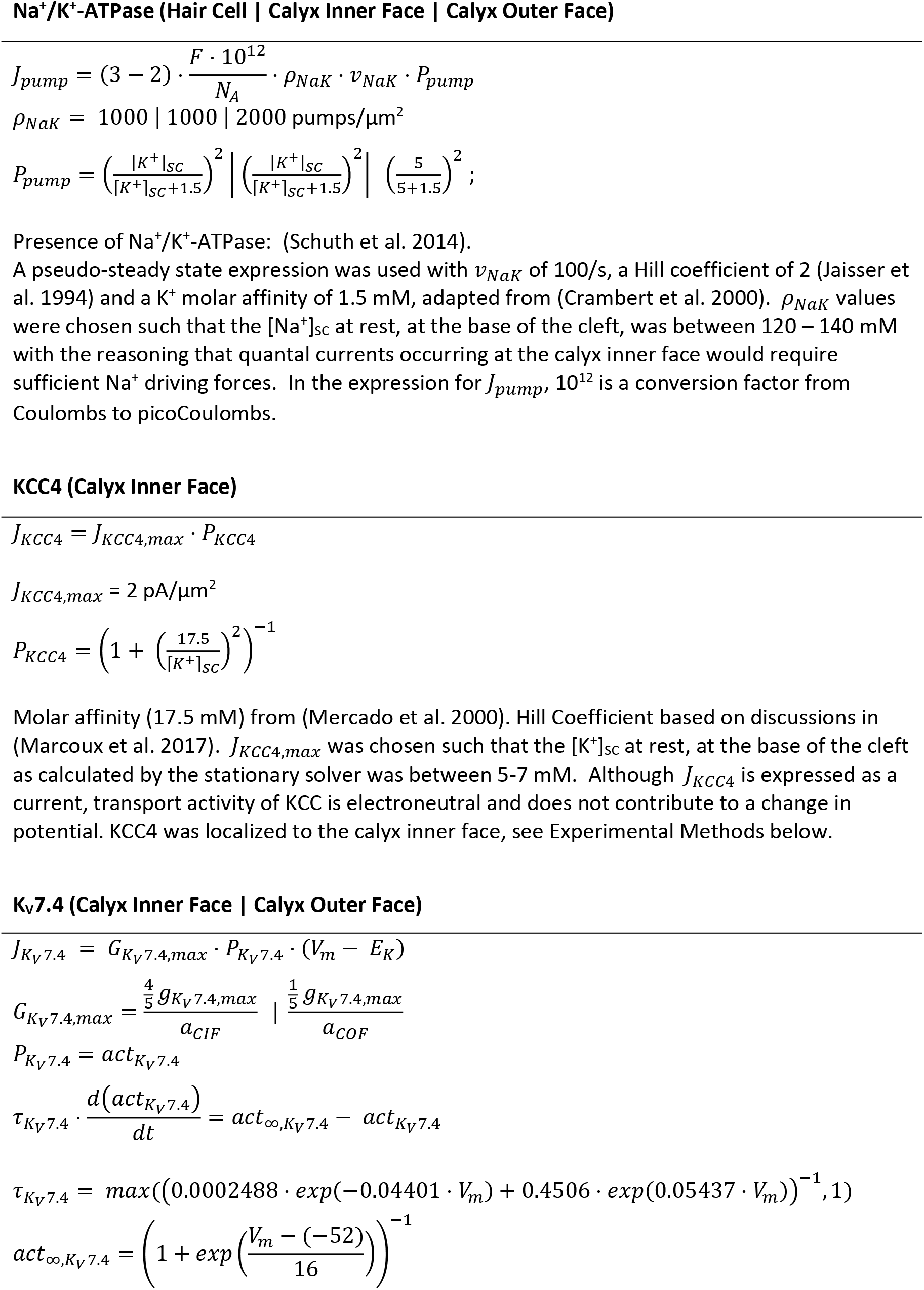
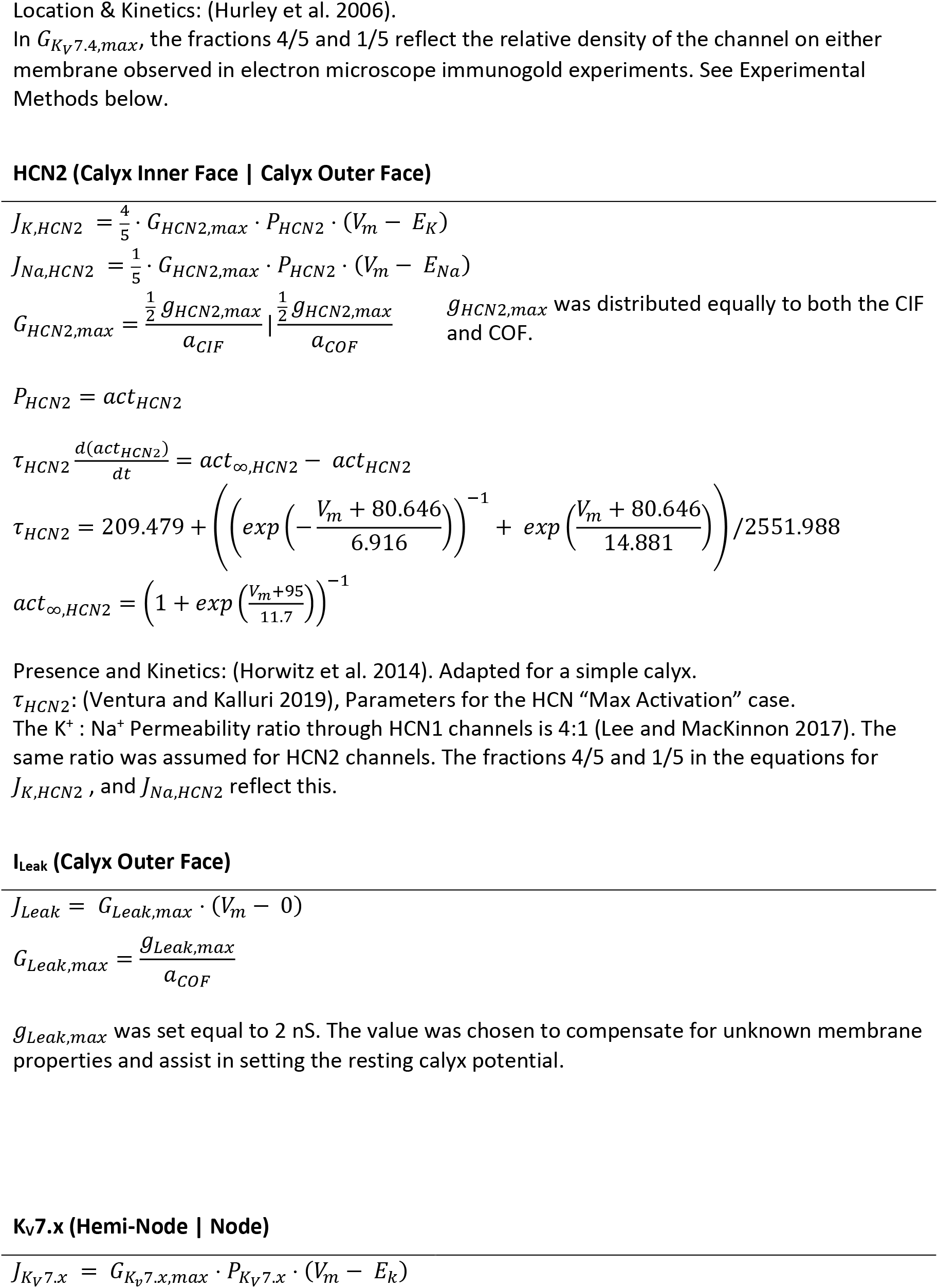
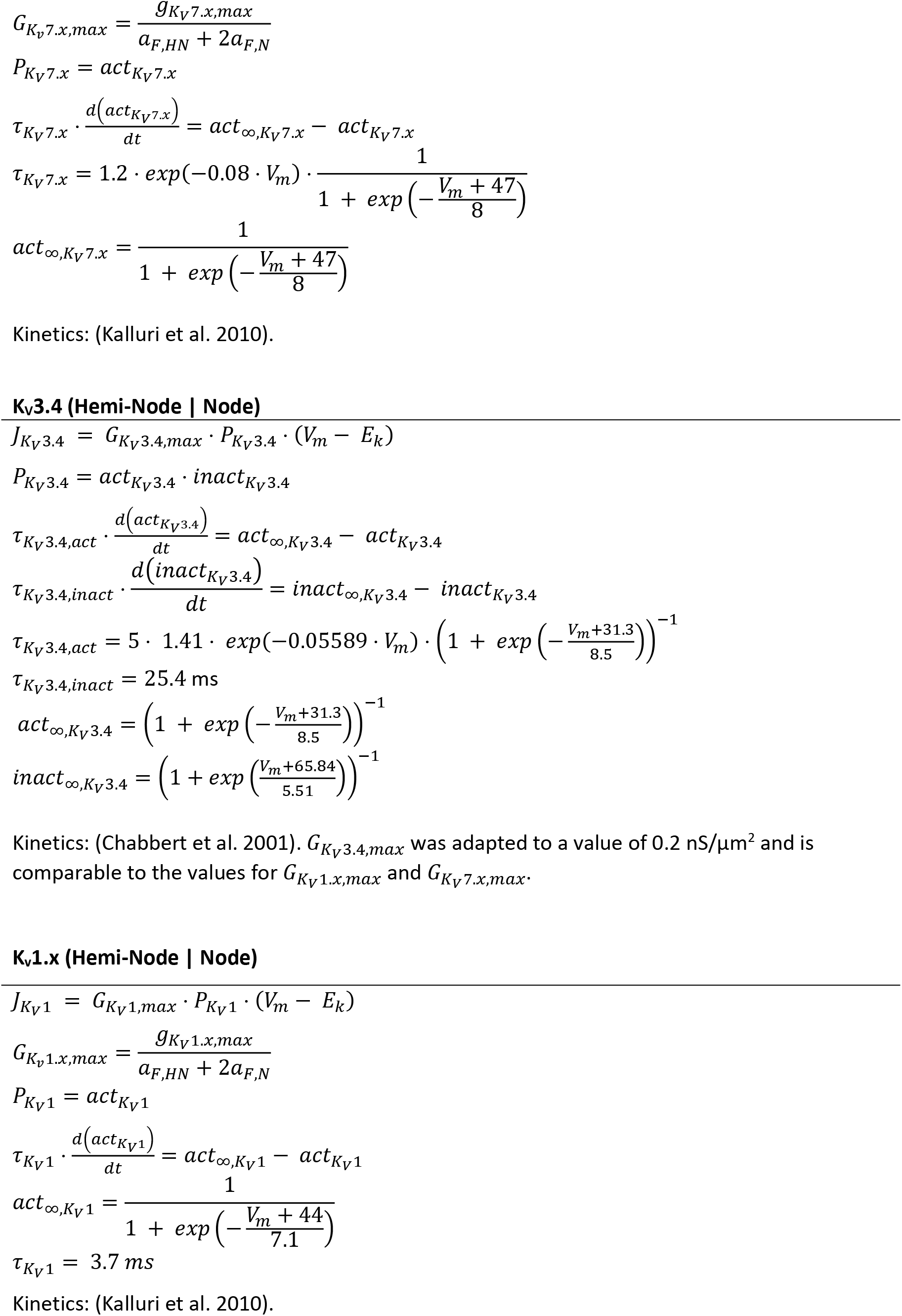
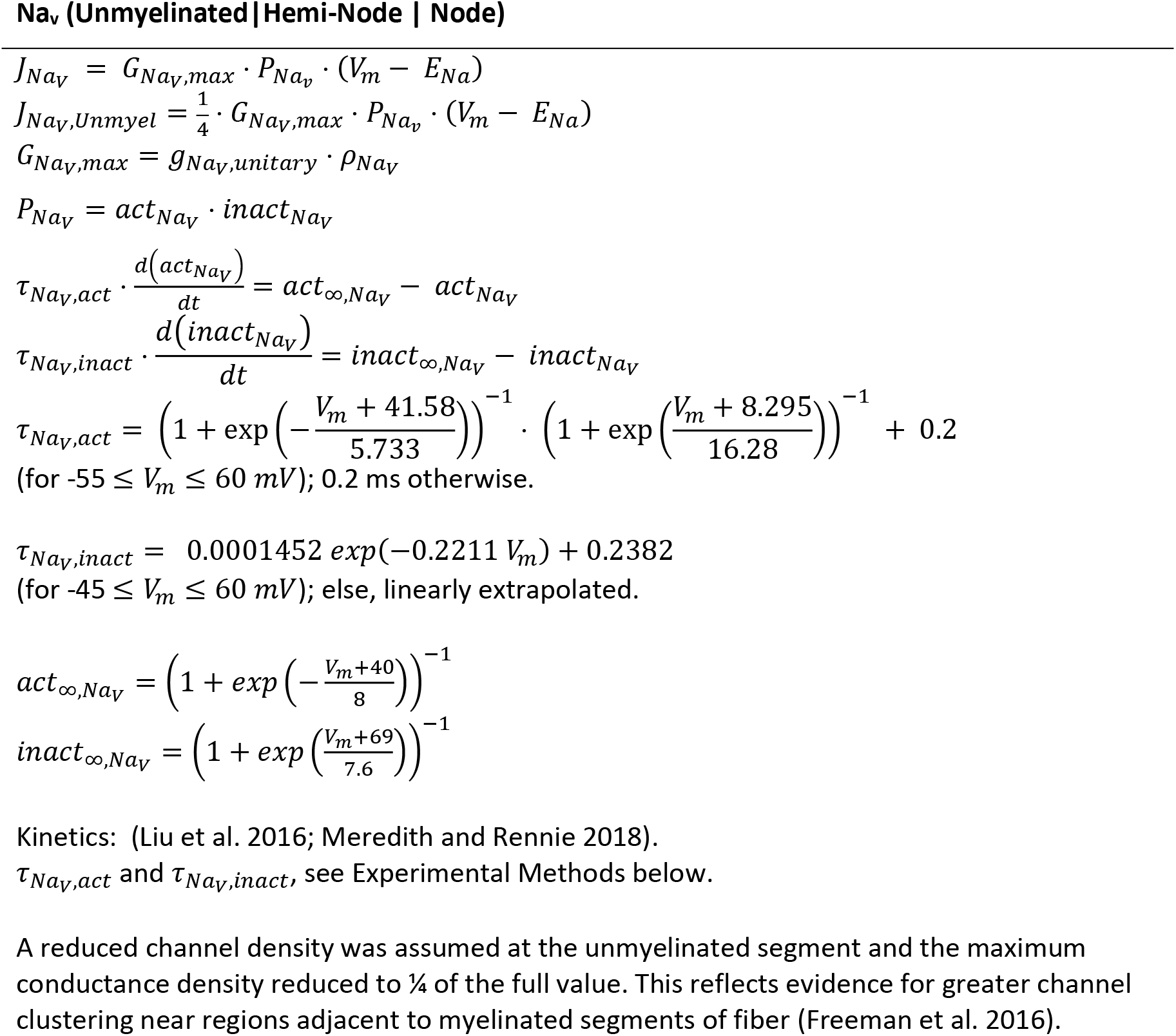
Currents Densities and Kinetics.

**Table S7.**
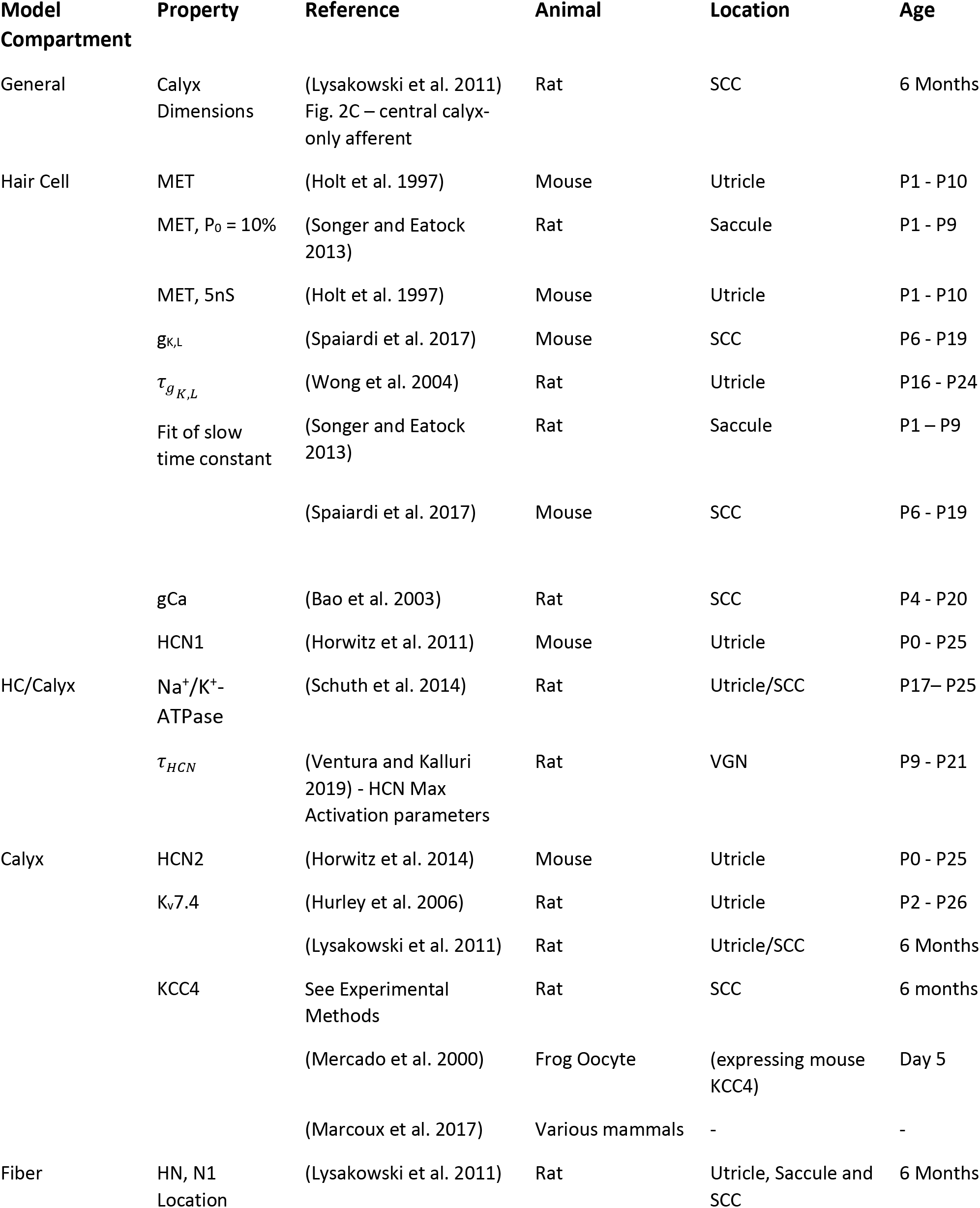
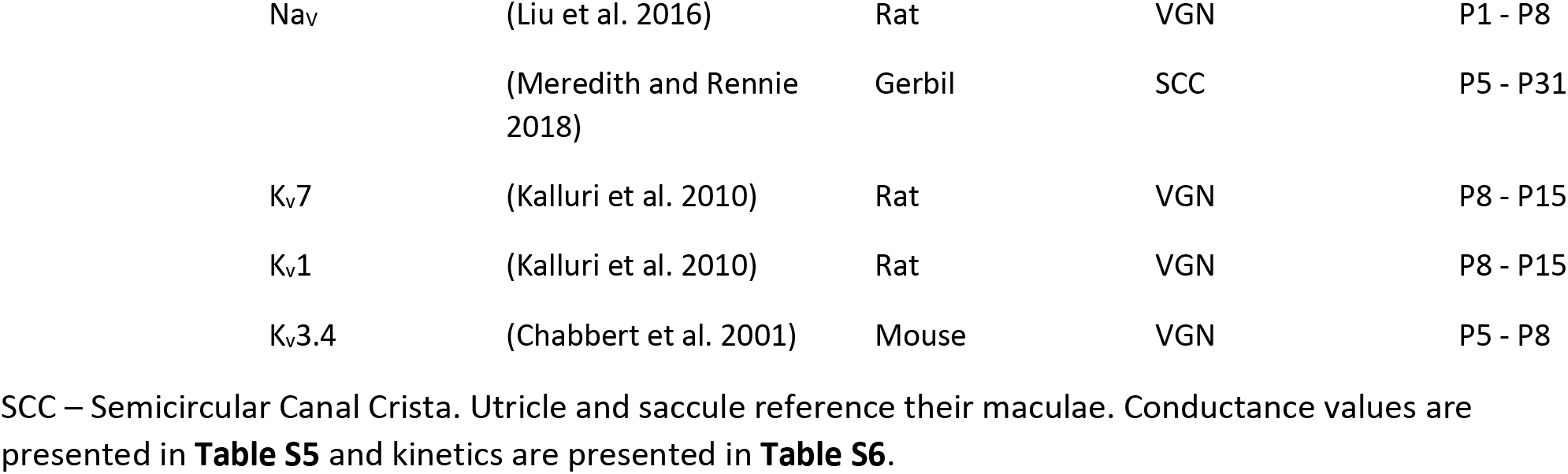
Sources.

## EXPERIMENTAL METHODS

Below we describe the experimental methods used to determine unpublished parameters listed in the Tables, and imaged cells shown in **Fig. 1A**.

### Na_V_ time constants (Table S6)

#### Whole-cell patch clamp recordings from mouse vestibular afferent neurons

The kinetics of postsynaptic (afferent) Na_V_ currents were obtained from mouse vestibular ganglion neurons isolated in the first postnatal month. Animals were handled in accordance with the *National Institutes of Health Guide for the Care and Use of Laboratory Animals*, and all procedures were approved by the animal care committee at the University of Chicago. Mice were deeply anesthetized with isoflurane and decapitated. Vestibular ganglia were dissected out and mechanically dissociated following enzymatic digestion (with 0.12% collagenase and 0.12% trypsin for 15-20 minutes at 37°C) and cultured overnight to remove satellite cells, in minimal essential medium with Glutamax (MEM, Invitrogen, Carlsbad, CA) supplemented with 10 mM HEPES, 5% fetal bovine serum (FBS), and 1% penicillin in 5% CO_2_-95% air at 37°C. Whole-cell patch clamp recordings were made with solutions designed to minimize contamination from K^+^ and Ca^2+^ currents. The external medium comprised (mM): 75 NaCl, 5.4 CsCl, 2.5 MgCl_2_, 75 TEACl, 5 HEPES, 10 D-glucose, titrated with CsOH to pH 7.4, with osmolality ~310 mmol/kg. The internal (pipette) solution contained: 148 CsCl, 0.8 CaCl, 3.5 Na_2_-creatine phosphate, 3.5 MgATP, 0.1 LiGTP, 5 EGTA, 5 HEPES, 0.1 Na-cAMP, titrated with CsOH to pH 7.4, and ~300 mmol/kg. Recordings were made at room temperature (23-25°C). Signals were delivered and recorded with a Multiclamp 700B amplifier, Digidata 1440A digitizer, and pClamp 10 or 11 software (Axon Instruments, Molecular Devices, Sunnyvale, CA), with low-pass Bessel filtering set at 10 kHz and sampling interval set at 5 μs. Seal resistance exceeded 1 GΩ. Membrane voltages were corrected off-line for a measured liquid junction potential of 6.4 mV and for series resistance errors. Analysis was performed with MATLAB (The MathWorks, Natick, MA), Clampfit (Axon Instruments, Molecular Devices) and Origin Pro 2018 (OriginLab, Northampton, MA). Series resistance (R_S_) ranged between 3 and 10 MΩ and was compensated by 75%. The holding potential was −66.4 mV. Activation and inactivation time courses were measured for currents evoked by depolarizing voltage steps following a pre-pulse to −127 mV to de-inactivate Na_V_ current. The currents were fit with the following Hodgkin-Huxley equations for Na_V_ channel activation (n=6) and inactivation (n=9):

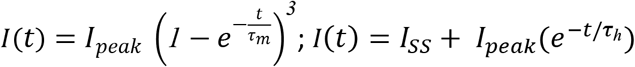

where *I* is current, *t* is time; *I_peak_* and *I_f_* are peak and steady-state (inactivated) currents, respectively; and *τ_m_* and *τ_h_* are the time constants for the activation and inactivation transitions, respectively.

#### Location of KCC, relative density of K_V_7.4 (Table S6) and image of hair cells and calyces (Fig. 1A): Confocal microscopy and immunocytochemistry

##### Fixation

Animals were handled in accordance with the *National Institutes of Health Guide for the Care and Use of Laboratory Animals,* and all procedures were approved by the IACUC at the Univ. of Illinois at Chicago. Long-Evans rats (Charles River Laboratories, Wilmington, MA) were deeply anesthetized with Nembutal (80 mg/kg), then perfused transcardially for 1 min with physiological saline containing heparin (2000 IU), followed by a dialdehyde fixative for 10 min (4% paraformaldehyde, 1% acrolein, 1% picric acid, and 5% sucrose in 0.1 M phosphate buffer (PB), pH 7.4). After decapitation, the head was post-fixed in the fixative for 20 min and then placed in 30% sucrose in 0.1 M PB. Vestibular and auditory organs were dissected out with the head immersed in PB. Decalcification of cochleas and removal of bone debris and otoliths from vestibular organs was accomplished through incubation in Cal-X solution. Background fluorescence was reduced by incubating the tissues in a 1% aqueous solution of sodium borohydride for 10 min. All organs were then placed in 30% sucrose in 0.1 M PB overnight.

##### Immunohistochemistry

Dissected tissues were gelatin-embedded and cut with a freezing sliding microtome. Tissue permeabilization was carried out with 4% Triton X-100 in PBS for 1 hr, and a blocking solution composed of 0.5% fish gelatin, 0.5% Triton X-100 and 1% BSA in 10 mM phosphate-buffered saline (PBS) was used for 1 hr. Samples were incubated in primary antibodies diluted 1:200 in blocking solution: goat anti-calretinin (AB1550), rabbit anti-KCC4 (AB3270) (all available or formerly available from Chemicon/Millipore-Sigma, St. Louis, MO) or rabbit anti-myosin 6 (a generous gift from Dr. Tama Hasson). The anti-calretinin antibody served as a marker of type II hair cells and calyx afferents (Desai et al. 2005). Specific labeling was revealed with these secondary antibodies (1:200 in the blocking solution): FITC-conjugated donkey anti-goat and rhodamine-conjugated donkey anti-rabbit (both from Chemicon, Temecula, CA). Sections were mounted on slides using a medium containing Mowiol (Calbiochem, Darmstadt, Germany) and examined on a confocal microscope (LSM 510 META, Carl Zeiss, Oberköchen, Germany). The relative density of Kv7.4 channels was based on electron microscopic (EM) immunogold experiments, with methods described in Hurley et al. (2006). In brief, 40-μm vibratome sections were incubated in primary antibody [1:200 rabbit anti-KCNQ4, a generous gift from Dr. Bechara Kachar] for 72 h and secondary antibody (1:50; colloidal gold-labeled goat anti-rabbit IgG; Amersham Biosciences, Piscataway, NJ) for 24 h. Colloidal gold staining was silver enhanced (4–8 min; IntenSE M kit; Amersham Biosciences). For more details, see Hurley et al. (2006).

